# Ethylene augments root hypoxia tolerance through amelioration of reactive oxygen species and growth cessation

**DOI:** 10.1101/2022.01.21.477196

**Authors:** Zeguang Liu, Sjon Hartman, Hans van Veen, Hongtao Zhang, Hendrika A.C.F. Leeggangers, Shanice Martopawiro, Femke Bosman, Florian de Deugd, Peng Su, Maureen Hummel, Tom Rankenberg, Kirsty L. Hassall, Julia Bailey-Serres, Frederica L. Theodoulou, Laurentius A.C.J. Voesenek, Rashmi Sasidharan

**Affiliations:** Plant Ecophysiology, Institute of Environmental Biology, Utrecht University, Padualaan 8, 3584 CH, Utrecht, The Netherlands; Department of Botany and Plant Biology, University of Geneva, 1211 Geneva, Switzerland; Plant Environmental Signalling and Development, Faculty of Biology, University of Freiburg, 79104 Freiburg, Germany; CIBSS – Centre for Integrative Biological Signalling Studies, University of Freiburg, 79104 Freiburg, Germany; Plant Sciences Department, Rothamsted Research, Harpenden, AL5 2JQ, UK; Computational and Analytical Sciences Department, Rothamsted Research, Harpenden, AL5 2JQ, UK; Botany and Plant Sciences Department and Center for Plant Cell Biology, University of California, Riverside, CA, 92521, USA

## Abstract

Flooded plants experience impaired gas diffusion underwater, leading to oxygen deprivation (hypoxia). The volatile plant hormone ethylene is rapidly trapped in submerged plant cells and is instrumental for enhanced hypoxia acclimation. However, the precise mechanisms underpinning ethylene-enhanced hypoxia survival remain unclear. We studied the effect of ethylene pre-treatment on hypoxia survival of primary *Arabidopsis thaliana* root tips. Both hypoxia itself and re-oxygenation following hypoxia are highly damaging to root tip cells and ethylene pre-treatments reduced this damage. Ethylene pre-treatment alone altered the abundance of transcripts and proteins involved in hypoxia responses, root growth, translation and reactive oxygen species (ROS) homeostasis. Through imaging and manipulating ROS abundance *in planta*, we demonstrate that ethylene limits excessive ROS formation during hypoxia and subsequent re-oxygenation and improves oxidative stress survival. In addition, we show that ethylene leads to rapid root growth cessation and this quiescence behaviour contributes to enhanced hypoxia tolerance. Collectively, our results show that the early flooding signal ethylene modulates a variety of processes that all contribute to hypoxia survival.

## Introduction

Anthropogenic climate change has increased global flood frequency and severity, challenging sustainable crop production (Hirabayashi et al., 2013; Voesenek and Bailey-Serres, 2015). Submerged plants experience a limitation of CO_2_ in photosynthesizing tissues, and oxygen (O_2_) deprivation (hypoxia) in respiring tissues, because of impaired gas diffusion underwater (Voesenek and Bailey-Serres, 2015). Especially in roots, characterized by much lower O_2_ levels (Vashisht et al., 2011; van Veen et al., 2016), mitochondrial respiration is arrested during hypoxia and glycolysis coupled to ethanolic fermentation becomes the main source of energy to fuel hypoxic cell survival (Loreti et al., 2016; Sasidharan et al., 2018). The severely reduced carbon fixation and high carbon demands of metabolic acclimation leads to severe energy reduction and carbon starvation. Paradoxically, de-submergence can further impair plant survival as returning to ambient light and O_2_ levels (normoxia) coincides with excessive formation of damaging reactive oxygen species (ROS) (Sasidharan et al., 2018; Yeung et al., 2018). Therefore, the rapid perception of submergence and timely activation of acclimation responses is essential for flooded plants to overcome hypoxia and re-oxygenation stress and increase chances of survival.

Flooding generates an array of endogenous signals that are perceived and processed by the plant to elicit adaptive responses (Sasidharan et al., 2018). However, accumulation of the gaseous plant hormone ethylene is considered the most robust signal for plant submergence detection (Voesenek and Sasidharan, 2013; Hartman et al., 2021). Indeed, ethylene biosynthesized during flooding is rapidly entrapped and perceived in submerged plant shoot and root tissues (Banga et al., 1996; Voesenek and Sasidharan, 2013; Hartman et al., 2019). This ethylene signal mediates a plethora of flood adaptive responses and facilitates acclimation to hypoxia in plants (van Veen et al., 2013; Sasidharan and Voesenek, 2015; Hartman et al., 2021). In addition, ethylene is essential for enhanced hypoxia survival and an augmented transcriptional induction of hypoxia responsive genes in multiple angiosperms (Peng et al., 2001; van Veen et al., 2013; Yamauchi et al., 2014; Hartman et al., 2020). Collectively, these reports show that the rapid accumulation of ethylene during flooding makes it an indispensable signal to mount a successful hypoxic acclimation response in plants.

Mimicking this ethylene accumulation by pre-treating plants with ethylene leads to enhanced hypoxia tolerance in *Rumex palustris* and several *Solanum* species and is associated with elevated hypoxia signaling (van Veen et al., 2013; Hartman et al., 2020). A similar response in *Arabidopsis thaliana* (Arabidopsis) allowed the identification of an important underlying molecular mechanism in this species (Hartman et al., 2019). Ethylene contributes to hypoxia anticipation and acclimation through enhancing the production and stabilisation of group VII Ethylene Response Factor transcription factors (ERFVIIs) prior to hypoxia (Hartman et al., 2019; Perata, 2020; Shi et al., 2020). The ERFVIIs are part of a mechanism that senses O_2_ and nitric oxide (NO) levels through the PROTEOLYSIS6 (PRT6) N-degron pathway of proteolysis and are the principal activators of the core transcriptional hypoxia response (Gibbs et al., 2011; Licausi et al., 2011; Gibbs et al., 2014; Bui et al., 2015; Gasch et al., 2016). Ethylene can stabilize ERFVIIs by scavenging NO through induction of non-symbiotic PHYTOGLOBIN1 (PGB1), already in ambient O_2_ (normoxia). This enhanced ERFVII stability consequently augments the core transcriptional hypoxia response when O_2_ levels diminish (Hartman et al., 2019; Hartman et al., 2021).

Interestingly, while this mechanism describes how ethylene accelerates the transcriptional core hypoxia response (Perata, 2020), quantitative differences in hypoxic gene induction can be poor predictors of tolerance (Loreti et al., 2016). Moreover, it currently remains unclear which genes, proteins and processes functionally contribute to enhanced hypoxia acclimation and survival. Finally, which of these processes are modulated by ethylene to increase survival during hypoxia also remains elusive. To investigate the processes underpinning ethylene-enhanced hypoxia tolerance, we assessed the transcriptome of Arabidopsis root tips after ethylene pre-treatment, subsequent hypoxia and re-oxygenation time points. We also quantified how the Arabidopsis proteome changes in response to an ethylene treatment. Our findings indicate that ethylene primarily promotes hypoxia tolerance through a collective response that includes enhanced hypoxia responses, improved ROS amelioration and cessation of root growth already prior to hypoxia.

## Results

### Ethylene enhances cell viability during both hypoxia and re-oxygenation

An ethylene pre-treatment enhances survival of subsequent hypoxia in Arabidopsis seedling root tips (Hartman et al., 2019). To uncover processes that contribute to this ethylene-enhanced hypoxic cell survival, we first investigated when root cells lose viability during hypoxia and re-oxygenation. The capacity of root tips to re-grow during hypoxia recovery (3 d) was used as a proxy for root meristem survival (Hartman et al., 2019). The results revealed that root tips of 4-day and 5-day old Arabidopsis Col-0 seedlings lose viability between 3.5 and 4.5 hours (h) of severe hypoxia, but that root tip survival is significantly prolonged after an ethylene pre-treatment (Figure 1A, Supplemental Figure S1A). Importantly, Evans Blue staining for cell membrane integrity revealed that a proportion of seedlings treated with 4 h of hypoxia do not lose root tip cell viability during the hypoxia period itself, but during the first hours of re-oxygenation (Figure 1B-C). Ethylene strongly reduced and delayed cell death during re-oxygenation (Figure 1B-C). When the hypoxia duration was prolonged (4.5h), cell death also occurred during hypoxia in addition to the re-oxygenation phase (Supplemental Figure S1B-C). Together, these results indicate that during shorter hypoxia stress (4 h in our experiments) root tips mainly die during the re-oxygenation phase, whereas longer hypoxia (4.5 h) results in meristem death in root tips during both the hypoxia and the subsequent re-oxygenation phase. Furthermore, ethylene enhances cell survival in root tips during both the hypoxia and re-oxygenation phase.

**Figure 1.**
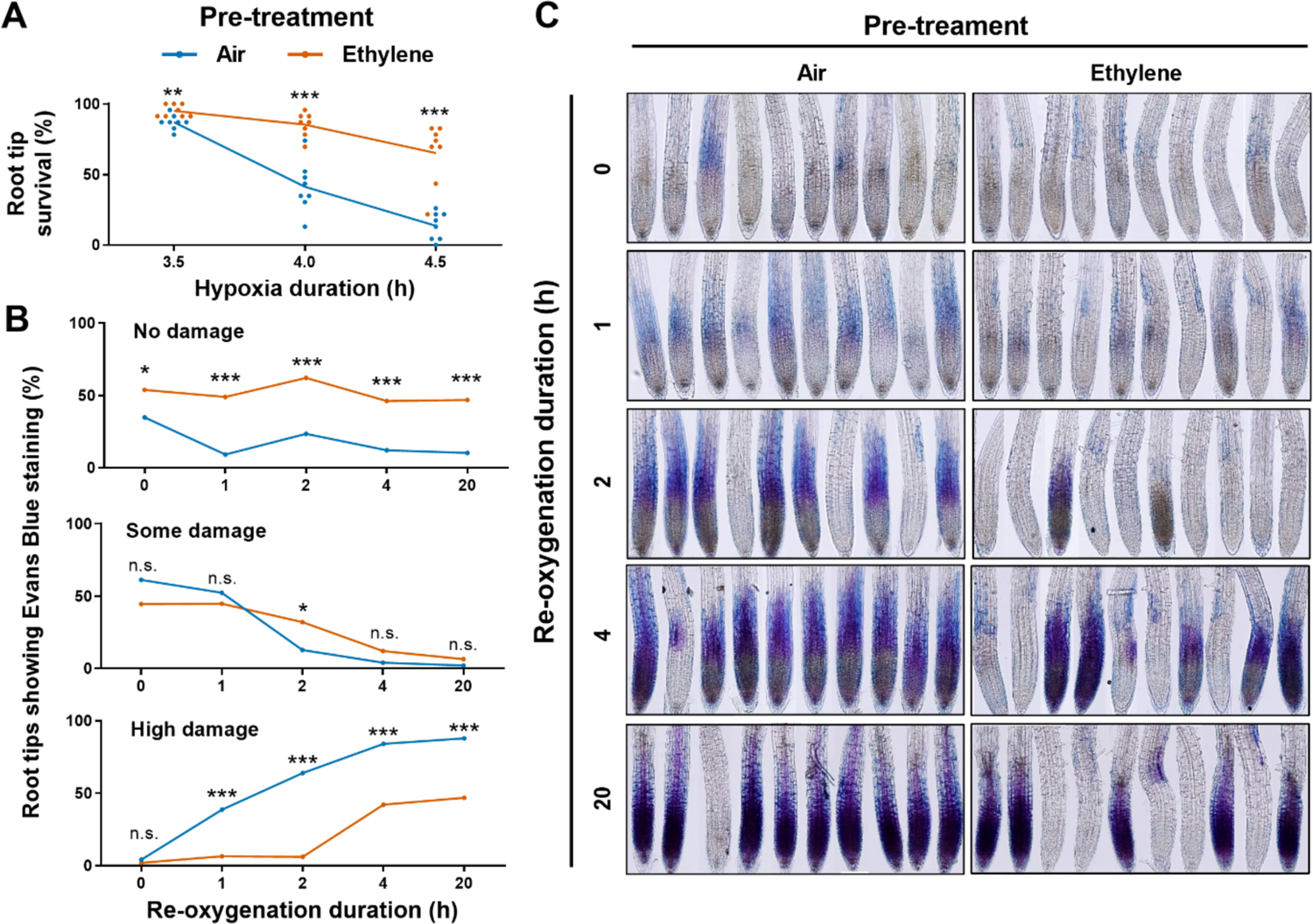
Ethylene pre-treatment improves cell viability during hypoxia and re-oxygenation. **(A)** Root tip survival of 4-day old Arabidopsis (Col-0) seedlings after 4 h of air (blue) or ∼5μL·L^-1^ ethylene (orange) treatment followed by hypoxia and 3 days of recovery (n=8 rows of 23 seedlings, Student’s t-test). **(B)** Classification and **(C)** visualization of Evans Blue (EB) staining for impaired cell membrane integrity in 4-day old seedling root tips after 4 h of pre-treatment with air or ∼5 μL·L^-1^ ethylene followed by 4 h hypoxia and re-oxygenation time points. Classification in (B), no damage = no EB staining, some damage = detectable EB staining, high damage = clear root-wide EB staining in elongation zone and root apical meristem, (C) scale bar = 100 µm. Asterisks indicate significant differences between air and ethylene per time point (n.s. - not significant, *p<0.05, **p<0.01, ***p<0.001, generalized linear model with a binomial error structure, n=44-54 root tips in B, n=10 random samples in C).

### Ethylene induces transcriptional changes that are maintained during subsequent hypoxia

To identify the processes and associated molecular identities associated with increased hypoxia tolerance after ethylene pre-treatment, microarray-based transcriptome profiling was performed (Supplemental Data, Sheet 1). The target tissues were root tips (0.5 cm) of 4-day old Arabidopsis Col-0 seedlings pre-treated either with 4 h of air or ethylene, and subsequently exposed to 2 and 4 h of hypoxia and 1 h of re-oxygenation (Figure 2A, Supplemental Figure S2, S3). Interestingly, whilst the number of Differentially Expressed Genes (DEGs) compared to the normoxic control continued to increase as the experiment progressed, the number of DEGs differing between air and ethylene (indicating an ethylene-specific effect) pre-treatment decreased over time (Figure 2B-C).

**Figure 2.**
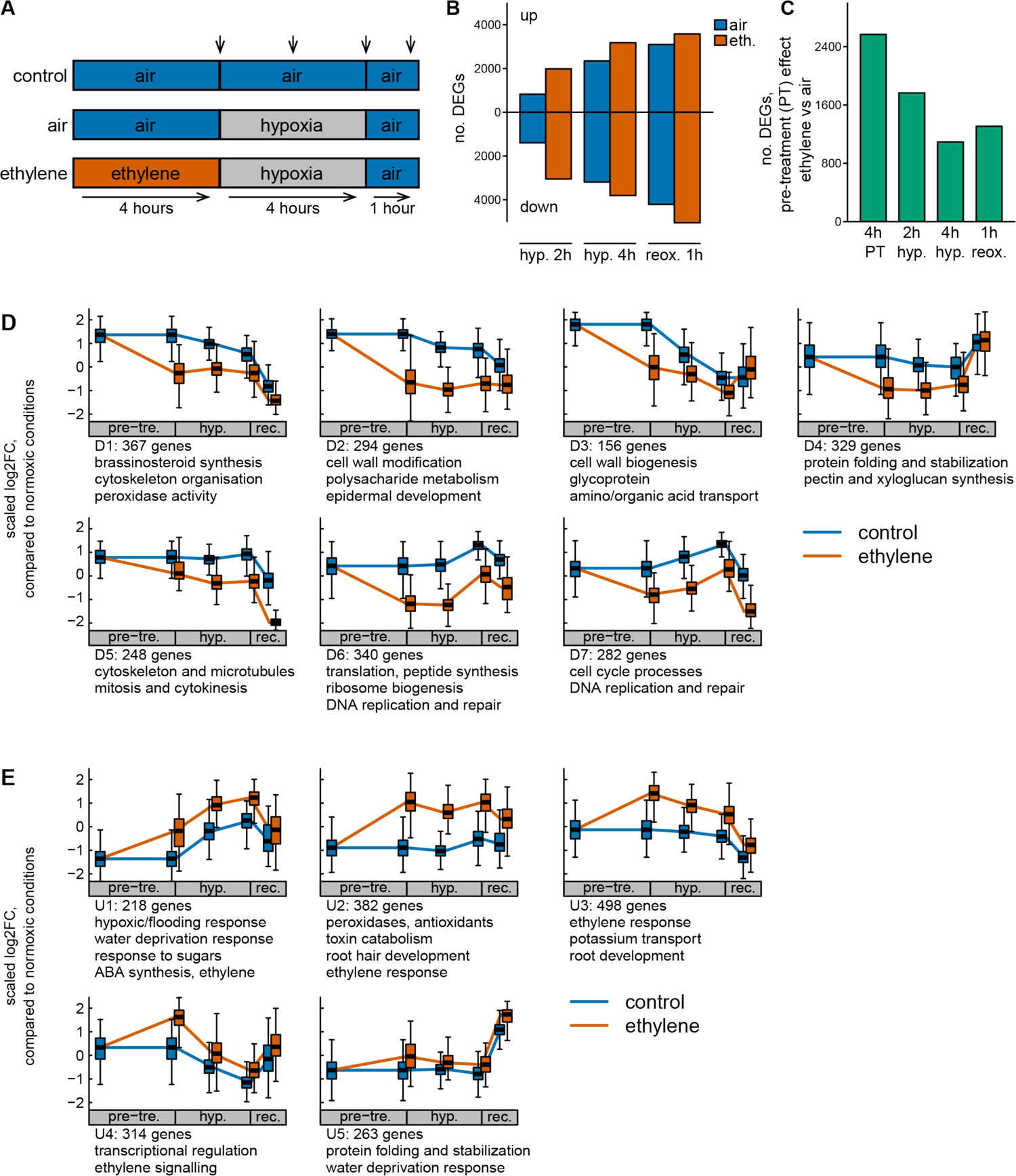
Ethylene leads to transcriptional reprogramming which is maintained during hypoxia and re-oxygenation. **(A)** Schematic overview of the experimental design, where arrows indicate the sampling timepoints. Arabidopsis (Col-0) 4-day old seedlings were pre-treated with air or ethylene (5μL·L^-1^) and subsequently exposed to 4 h hypoxia (O_2_ levels reach 0% after ∼40 minutes after the start of treatment) and 1 h of normoxia in the dark. Approximately 184 root tips (<0.5 mm from tip) were harvested per experimental replicate (n=3) after the pre-treatment, 2 h and 4 h of hypoxia and 1 h of re-oxygenation. **(B)** The number of differentially expressed genes (DEGs, p < 0.001) relative to the controls (air-normoxia) of the same timepoint for both air and ethylene pre-treatment. **(C)** The number of DEGs (p<0.001) between air and ethylene pretreated plants, for each harvest time point. **(D)** and **(E)** Clusters from hierarchical clustering. Key processes and functions enriched in each cluster based on gene ontology (GO) enrichment. Full lists of DEGs per cluster and associated GO terms are available in the Supplemental Data (sheet 1-4). Shown FCs are relative to the normoxic control of the corresponding timepoints, and were mean and standard deviation scaled by gene.

Genes that behaved differently depending on the ethylene pre-treatment were of primary interest to explain differences in ethylene-mediated hypoxia and re-oxygenation tolerance. Ethylene-mediated differences in expression were maximal directly after the pre-treatment (Figure 1C). Of the genes that differed during hypoxia treatment, almost all were also already differentially expressed directly after the ethylene pre-treatment (Figure 2C-E). To investigate how these early ethylene effects on the transcriptome developed throughout the treatment, we grouped the ethylene pre-treatment dependent genes (Differentially Regulated Genes, DRGs) into 12 unique expression profiles by hierarchical clustering (Figure 2D, E).

Seven distinct profiles were associated with a strong downregulation of transcripts by ethylene that either strengthened or counteracted responses to hypoxia and re-oxygenation (D1-7, Figure 2D). A reverse pattern, where clusters were enhanced by ethylene, was found in 5 profiles (U1-5, Figure 2E). In none of these clusters were the pre-treatment differences further increased or decreased during hypoxia. Though these DEGs were mostly maintained during hypoxia, the difference between air and ethylene pre-treatment typically declined over time (Figure 2D-E). We conclude that the induction of ethylene signaling prior to hypoxia induces a major transcriptome reconfiguration in root tips and that most of these changes are maintained during subsequent hypoxia, and to a lesser extent during re-oxygenation.

We also identified hypoxia- and re-oxygenation-responsive processes that were not modulated by the ethylene pre-treatment (Supplemental Figure S5). The hypoxic induction of transcripts that continued during the reoxygenation, were associated with protein folding and hypoxia, heat, and ROS responses. Genes that were continuously downregulated during hypoxia and re-oxygenation were associated with ion transport and root development. These genes may be important for acclimation to hypoxia and re-oxygenation but are unlikely to be involved in ethylene-mediated hypoxia tolerance. To better understand the influence of ethylene on transcriptional reprogramming, we went on to consider individual DRGs.

### Ethylene elicits a transcriptional signature associated with root growth cessation, ROS homeostasis and stress responses

The nature of the genes present in each cluster (Figure 2D, E) was assessed by gene ontology (GO) enrichment analysis. Widespread ethylene signaling-related GO terms were detected in the ethylene-enriched clusters (U1-U4; Figure 2E). Additional ethylene-enhanced GO terms included hypoxic/flooding responses, response to sugars, ABA biosynthesis (U1), toxin catabolism, antioxidants (U2) and root development (U3). Ethylene-downregulated GO terms were indicative of a general reduction in root growth, development, and cellular maintenance, including brassinosteroid biosynthesis (D1), cell wall biogenesis (D3), mitosis and cytokinesis (D5), translation and ribosome biogenesis (D6), and cell cycle (D7).

Ethylene is an established inhibitor of root growth and development (Le et al., 2001), and both known and novel ethylene-driven gene clusters and genes indicative of ethylene-mediated growth cessation were identified. Ethylene suppresses root cell enlargement in the elongation zone and the epidermis through auxin signaling (Růžička et al., 2007; Swarup et al., 2007; Vaseva et al., 2018). Accordingly, cluster U3 contained known ethylene-mediated players in auxin upregulation and transport, including *ANTHRANILATE SYNTHASE α1* (*ASA1*), *ERF1* and *HOMEOBOX52* (*HB52*) (Mao et al., 2016; Miao et al., 2018). Cluster D2 was enriched in genes associated with cell wall modification being suppressed by ethylene. Apart from limiting growth by elongation, ethylene is also reported to reduce cell proliferation (Street et al., 2015). Accordingly, ethylene pre-treatment led to downregulation of mitosis, DNA replication and cyclin genes in cluster D5, D6, D7.

In addition to known ethylene-mediated growth suppressing pathways, our results revealed effects on growth regulators not previously associated with ethylene signaling. The development of the root relies heavily on the PLETHORA (PLT) 1 and 2 transcription factors (Aida et al., 2004; Galinha et al., 2007), the GRAS family proteins SCARECROW (SCR) and SHORTROOT (SHR) (Sozzani et al., 2010), TEOSINTE-BRANCHED CYCLOIDEA (TCP) 20/21 (Shimotohno et al., 2018) and SHORT HYPCOTYL (SHY) 2 (Tian and Reed, 1999). Notably, these growth stimulating genes, *PLT1* (D7), *PLT2* (D5), *SCR* (D1), *SHR* (D3), *TCP21* (D2) and *SHY2* (D6) were all downregulated after ethylene treatment. Genes associated with biosynthesis of the root growth stimulating hormone brassinosteroid (D1) were also downregulated by ethylene (Müssig et al., 2003). Moreover, genes associated with biosynthesis of the root growth suppressor ABA were induced (U1) alongside many SnRK2s (2.6 U1; 2.7 U3; 2.9 U4; 2.10 U1) that mediate ABA and stress-induced root system architectural changes (McLoughlin et al., 2012; Kawa et al., 2020). Together these results indicate that ethylene leads to both known and novel transcriptional changes that are indicative of growth cessation in the root tip, which are maintained during hypoxia and re-oxygenation.

Ethylene also caused differential regulation of genes associated with ROS homeostasis and antioxidant activity that modulate root growth and oxidative stress responses. (Figure 2C, D; D1, U2) (Supplemental Figure S6). These included *UPBEAT1*, a transcription factor that stimulates cell proliferation and root growth by diminishing superoxide levels through the suppression of peroxidases (Tsukagoshi et al., 2010) and previously linked to waterlogging tolerance in *Brassica napus* through growth repression (Guo et al., 2020). *UPBEAT1* was ethylene induced (U4), whilst its direct target peroxidase was ethylene suppressed (D1). Ethylene downregulated *MYB30* (D1), a ROS-inducible root growth stimulating transcription factor that stimulates cell proliferation (Mabuchi et al., 2018). Collectively, this behavior of key transcription factors and genes, controlling or mediated by ROS, suggest that root growth could be reduced by ethylene through modulating ROS production (D1) and scavenging (U2).

Ethylene also controls many stress related genes, culminating in cluster U1, where an initial transcriptional induction by ethylene was further continued during hypoxia. These included a range of genes classically associated with hypoxia, such as *ALCOHOL DEHYDROGENASE 1* (*ADH1*) and *PGB1*. Two *ERF-VII*s, *HYPOXIA RESPONSIVE ERF 1* (*HRE1*) and *RAP2.3,* were also ethylene induced (U2). *CBL-INTERACTING PROTEIN KINASE 25*, recently implicated in root hypoxia tolerance (Tagliani et al., 2020) was also strongly upregulated by ethylene (U1). Similarly, transcripts of *HYPOXIA UNKOWN PROTEIN 54* (AT4G27450) were already enhanced in response to ethylene. Ethylene also enhanced genes associated with water deprivation responses, which is a typical stress during re-oxygenation (Yeung et al., 2019). Apart from ABA biosynthesis and signaling (U1) this included ROS amelioration (U2) consisting of several peroxidases, including *ASCORBATE PEROXIDASE2* (Figure 2E, S6). However, several peroxidases were also enriched in the ethylene downregulated cluster D1.

Collectively, this transcriptomic analysis demonstrates that ethylene mediates the majority of DEGs already prior to hypoxia stress and controls processes that include root growth cessation, hypoxia responses and ROS homeostasis and amelioration.

### Ethylene enhances proteins involved in ROS homeostasis and both mitochondrial and anaerobic respiration

In addition to transcription, changes in translational activity and proteolysis are also considered to be vital for protein homeostasis and therefore stress acclimation responses (Chang et al., 2000; Juntawong et al., 2014; Millar et al., 2019). *De novo* protein synthesis is dramatically impaired during severe hypoxia (Sachs et al., 1980) and the capacity to produce a new set of proteins is essential for hypoxia acclimation (Ellis et al., 1999). Providing mild hypoxia allows *de novo* protein synthesis and is considered crucial for successful acclimation to severe anoxia (Chang et al., 2000). We therefore profiled the proteome of the root tips after four hours of pre-treatment with ethylene or air, the timepoint at which most ethylene-mediated transcriptional differences were established and the plants would still display strong translation capacity (Figure 2B).

With a cut-off of 2 or more detected peptides for protein recognition, we quantified 6525 Arabidopsis proteins using isobaric multiplex tandem mass tag (TMT™)-based quantitative mass spectrometry (MS) (Supplemental Data, Sheet 9). Ethylene enhanced the abundance of 435 proteins and decreased the abundance of 350 proteins (Figure 3A). A GO analysis revealed an enrichment of processes that to some extent mirrored those found in the transcriptome analysis (Figure 3B). This included an upregulation of water deprivation responses, protein folding and anaerobic metabolism, and a downregulation of DNA replication and translation. Additionally, we found an auxin signature, indicating alterations in the root developmental program. GO terms that were unique to the proteomics dataset were associated with the mitochondria, lipid metabolism, and multivesicular bodies. Interestingly, a direct comparison of transcript and protein responses revealed little correlation (Figure 3C). Many ethylene-regulated proteins did not change at the transcript level, with the important exceptions of PGB1, HUP36, AT1G21400 (THDP-binding superfamily protein), and Nitrate Reductase 1 (Figure 3C--D). Among up regulated proteins there was only a mild but significant enrichment of genes with a corresponding regulation at the transcript level (odds-ratio 95% CI =1.8-3.2, P < 0.001), and this was even less pronounced for down-regulated proteins (odds-ratio 95% CI =1.1-2.0, P < 0.05).

**Figure 3.**
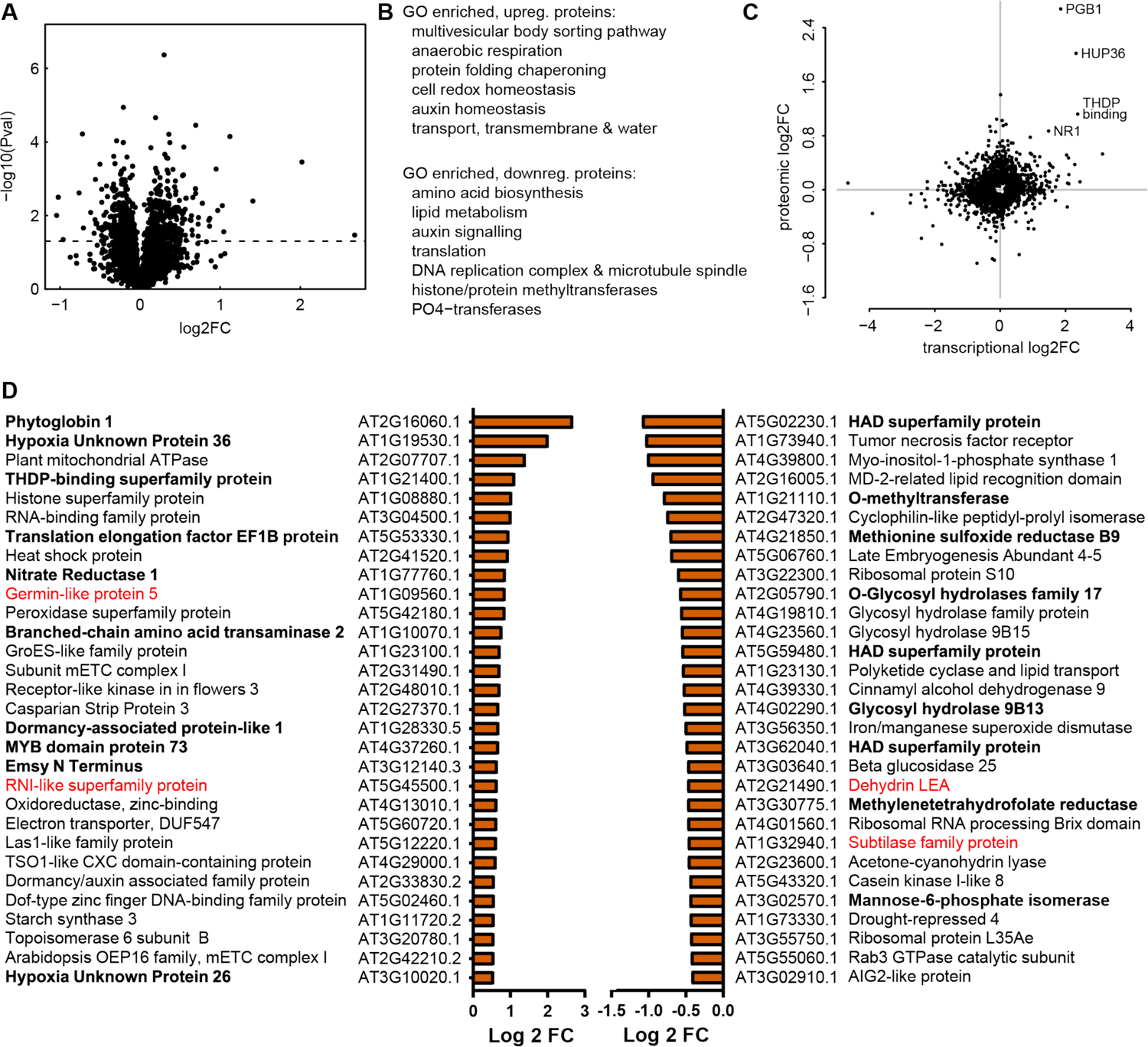
Ethylene treatment modulates the Arabidopsis root tip proteome. **(A)** Volcano plots showing the quantitative change in protein abundance upon 4 h of ethylene treatment (5 μL·L^-1^; the pre-treatment) and the statistical significance. **(B)** Key processes, functions and locations enriched among up and down regulated proteins based on gene ontology (GO) enrichment. Full lists of GO terms are available in the Supplemental Data (Sheet 10). **(C)** Direct comparison of transcriptomic and proteomic response for individual genes. **(D)** The 30 strongest up and down regulated proteins are shown based on fold change. Proteins in bold are also transcriptionally regulated in the corresponding direction. Proteins in red showed an opposite transcriptional response. All proteins shown are significantly different (p < 0.05, Student’s test, n=5).

Protein levels can be modulated by changes in translation and proteolysis. The most differentially ethylene-enriched protein detected was PGB1. Although *PGB1* is transcriptionally induced by ethylene, the NO scavenging PGB1 protein influences protein abundance by limiting NO-dependent proteolysis of Met_1_Cys_2_(MC)-initiating ERFVII proteins through the PRT6 N-degron pathway (Gibbs et al., 2014; Gibbs et al., 2018; Hartman et al., 2019). In the current dataset, 29 other MC-initiating proteins were detected, but none were enriched in response to ethylene (Supplemental Table S1). However, ethylene treatment reduced the abundance of three MC-proteins with unknown biological function (AT5G12850, AT2G10450 and AT2G26470; Supplemental Table S1). A comparison with previously reported proteins up-regulated in both the PRT6 N-degron loss-of-function mutants *prt6* and *ate1ate2* showed that at least 3 proteins (PGB1, HUP54 and AT5G63550) were also induced by ethylene (Supplemental Table S2, (Zhang et al., 2015)). In general, the proteome profiling results suggest that ethylene regulates protein abundance partially through transcription, but also through yet unidentified changes in translation and proteolysis.

Notable findings among the top 30 most upregulated proteins were PGB1, HUP26 and HUP36 (Figure 3D). PGB1 and HUPs are core hypoxia responsive genes (Mustroph et al., 2010), typically associated with hypoxia acclimation. Interestingly, they were enhanced in response to ethylene despite the absence of hypoxia. To assess whether this ethylene-enhanced abundance of HUP proteins is beneficial for surviving subsequent hypoxia, we evaluated root tip hypoxia survival in overexpression lines of the three HUPs (−26, −36 and −54) upregulated in the proteomics dataset (Supplemental Figure S7). The results indicated that, compared to wild-type Col-0, overexpression of *HUP26* and *36,* but not *HUP54* resulted in a higher basal hypoxia tolerance even in the absence of an ethylene pre-treatment. Additionally, *HUP36* and *HUP54* overexpression increased ethylene-induced hypoxia tolerance relative to wild-type (Supplemental Figure S7). This provides support for the involvement of HUP26 and HUP36 in ethylene-mediated hypoxia acclimation of root tips.

Amongst the top induced proteins were also several proteins associated with mitochondrial respiration, located in the mitochondrial electron transport chain (mETC), NITRATE REDUCTASE1 (NR1), and peroxidases. Among the top downregulated proteins were several ribosomal proteins. These findings echoed the GO term enrichment amongst the ethylene regulated proteins (Figure 3B, C) and the GO terms identified at the transcriptional level. In summary, these results reveal that an ethylene treatment quantitatively alters the Arabidopsis root tip proteome and pinpoints modulation of mitochondria, ROS-redox and amelioration, anaerobic metabolism and translation as key processes that are affected and could aid subsequent hypoxia acclimation.

### Ethylene improves antioxidant levels and ROS amelioration during hypoxia and re-oxygenation

Both the transcriptomics and proteomics data suggest that ethylene not only mediates ROS homeostasis and antioxidant activity during the pre-treatment but also during subsequent hypoxia and re-oxygenation. As ROS signaling is essential for flooding acclimation, and re-oxygenation is associated with toxic levels of ROS accumulation (Yeung et al., 2019), we explored the role of ethylene in ROS homeostasis. Ethylene affected the transcript and protein abundance of several peroxidases (Figure 2-3, Supplemental Figure S6). Considering this and given that the peroxidase-mediated control of hydrogen peroxide (H_2_O_2_) strongly contributes to cellular antioxidant capacity (Das and Roychoudhury, 2014), we assessed total soluble peroxidase activity in Arabidopsis root tips. We found higher peroxidase activity in the ethylene pre-treated samples both during the ethylene pre-treatment and subsequent hypoxia and re-oxygenation phase (Figure 4A). The Glutathione-Ascorbic acid (AA) pathway (Foyer and Noctor, 2011) also plays a major role in ROS homeostasis and changes in glutathione and AA are established indicators of changes in cellular redox status. Therefore, we assessed how ethylene modulates the levels of these compounds. No differences were observed in AA content in response to ethylene or subsequent hypoxia and re-oxygenation (Figure 4B). However, our results revealed that ethylene may alter the antioxidant capacity through total glutathione abundance (Figure 4C, D). After an ethylene pre-treatment, both the reduced (GSH) and oxidized glutathione (GSSG) levels increased, and this effect was maintained after 4 hours of hypoxia, but not during re-oxygenation (Figure 4C, D). However, the GSH:GSSG ratio did not change during any of the treatments (Figure 4C-E). These results are consistent with the report that ethylene insensitive mutants are impaired in glutathione biosynthesis, but not in AA production during abiotic stress (Yoshida et al., 2009). Together, these results reveal that ethylene may alter ROS homeostasis through changes in at least peroxidase activity and glutathione content.

**Figure 4.**
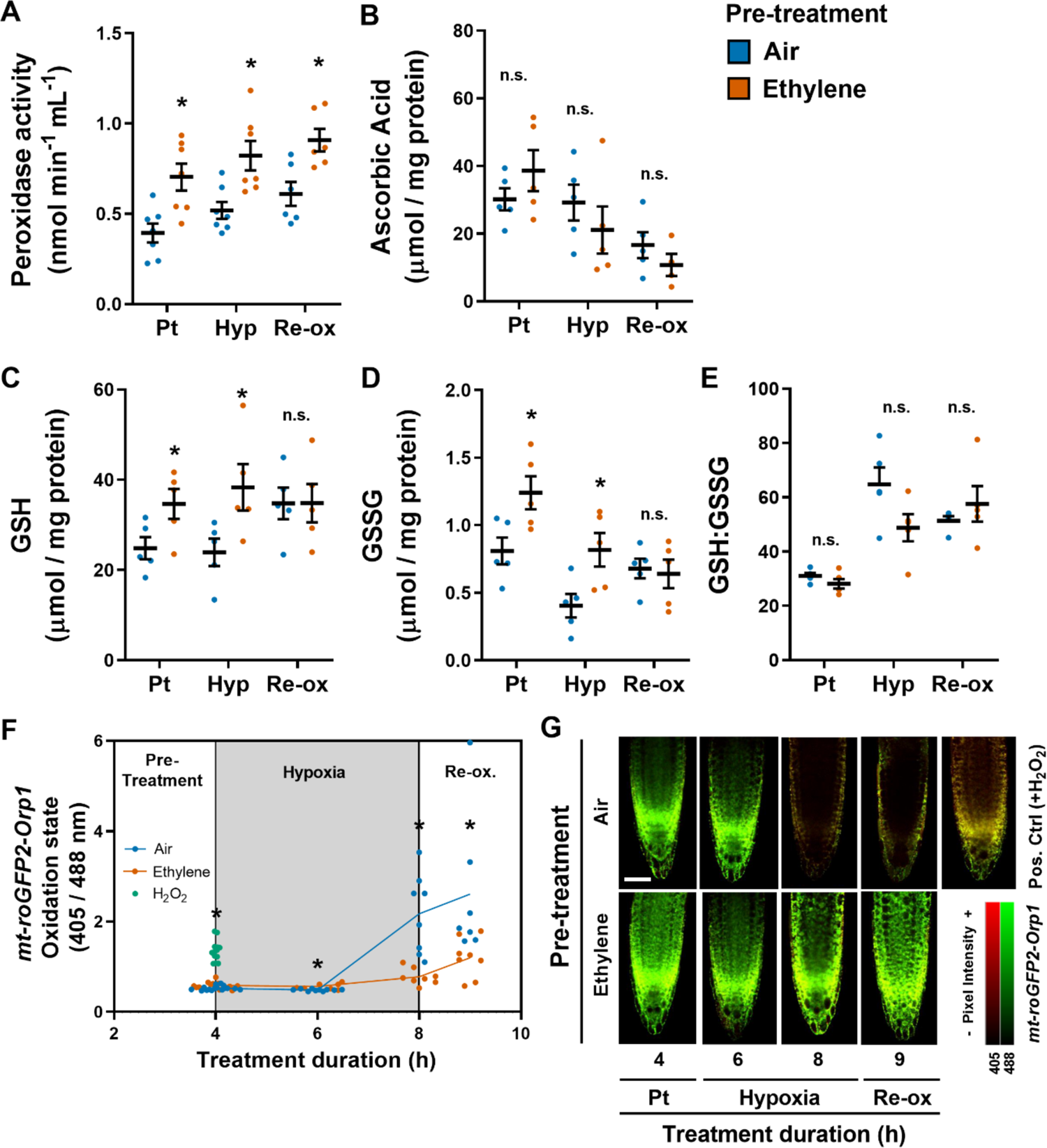
Ethylene mediates antioxidant capacity and ROS homeostasis after pre-treatment and subsequent hypoxia and re-oxygenation. (A-E) Peroxidase activity (A), ascorbic acid (B), reduced glutathione (GSH, C), oxidized glutathione (GSSG, D) and the GSH:GSSG ratio (E) in 5-day old Arabidopsis Col-0 root tips after 4 h of air (blue) or ∼5 μL·L^-1^ ethylene (orange) pre-treatment (Pt) followed by 4 h of hypoxia (Hyp) and 1 h of re-oxygenation (Re-ox) (n=5-6). (F, G) Quantification (F) and representative overlay images (G) of the oxidation state of mitochondrial targeted roGFP2-Orp1 in Arabidopsis root tips after 4 h of pre-treatment with air (blue) or ∼5μL·L^-1^ ethylene (orange) followed by 2 and 4 h of hypoxia and 1 h of re-oxygenation. The oxidation state of roGFP2-Orp1 is calculated from 405nm/488 excitation ratios, where a higher oxidation rate corresponds to elevated H_2_O_2_ in mitochondrial matrix (Nietzel et al., 2018). A positive control was included at the 4 h time-point by adding 20mM H_2_O_2_ to the root tip, 30 minutes prior to imaging (green). Asterisks indicate significant differences between air and ethylene per time point (*p<0.05, Student’s t-test, n=7-16 root tips).

Next, we assessed if ethylene pre-treatment altered ROS levels during subsequent hypoxia and re-oxygenation. For this, we used two different methods to visualize H_2_O_2_ in Arabidopsis root tips. Firstly, we used the transgenic H_2_O_2_-reporter line roGFP2-Orp1 which allows *in vivo* visualization and quantification of H_2_O_2_ in specific organelles at a subcellular level (Nietzel et al., 2019). Since a large portion of ROS is formed inside mitochondria through excessive electron flow through the ETC (Vanlerberghe, 2013) and ethylene led to differences in several proteins located in the mETC (Figure 3B, C), we decided to study the mitochondrion-specific (mt-) roGFP2-Orp1(Nietzel et al., 2019). Application of H_2_O_2_ to root tips prior to imaging, functioned as a positive control and showed an enhanced roGFP2-Orp1 oxidation state, indicative of higher H_2_O_2_ levels (Figure 4F, G). Ethylene pre-treatment alone slightly increased mitochondrial H_2_O_2_ levels in Arabidopsis root tips (Figure 4F, G). After both 4 h of hypoxia and 1 h of re-oxygenation, H_2_O_2_ levels increased drastically in air pre-treated root tips. However, excessive ROS formation observed in the mitochondria was strongly limited after an ethylene pre-treatment. As a second approach to visualize H_2_O_2_ we used 3,3’-Diaminobenzidine (DAB) staining. Again, an ethylene pre-treatment limited root tip H_2_O_2_ accumulation during subsequent hypoxia and re-oxygenation timepoints (Supplemental Figure S8). Collectively, these results suggest that ethylene is essential to boost antioxidant capacity and limit the excessive ROS formation during subsequent hypoxia and re-oxygenation in Arabidopsis root tips.

### Ethylene improves oxidative stress tolerance

Our results show that a large proportion of Arabidopsis root tips lose cell integrity during re-oxygenation (Figure 1B, C, Supplemental Figure S1B, C). We therefore wondered whether the pronounced increase in H_2_O_2_ during hypoxia and re-oxygenation (Figure 4F, G) leads to oxidative stress and subsequent reduced survival in these root tips. Or conversely, that impaired cell integrity leads to elevated H_2_O_2_ levels as ROS homeostasis can no longer be maintained. To untangle this question, we first examined whether ethylene-mediated ROS amelioration contributes to enhanced hypoxia survival by applying the pharmaceutical ROS scavenger potassium iodide (KI). Indeed, like an ethylene pre-treatment, the application of either 1- or 5-mM KI strongly boosted hypoxia root tip survival (Figure 5A). While these results demonstrate that improved ROS amelioration contributes to hypoxia survival, it does not reveal a direct effect of ethylene on enhanced oxidative stress tolerance. Therefore, we tested whether an ethylene pre-treatment could improve oxidative stress survival independent of hypoxia stress. Accordingly, small droplets of H_2_O_2_ were applied directly to the root tips after an air or ethylene pre-treatment to induce oxidative stress. Application of 6 mM H_2_O_2_ strongly reduced root tip survival, but this impaired survival could be rescued by an ethylene pre-treatment (Figure 5B). This ethylene-mediated ROS tolerance was dependent on the established ethylene-signaling EIN3/EIL1 transcription factor complex (Alonso et al., 2003), as the *ein3-1eil1-1* double mutant no longer benefited from an ethylene pre-treatment (Figure 5B). Taken together, these results suggest that ethylene-mediated ROS amelioration contributes to hypoxia tolerance in Arabidopsis.

**Figure 5.**
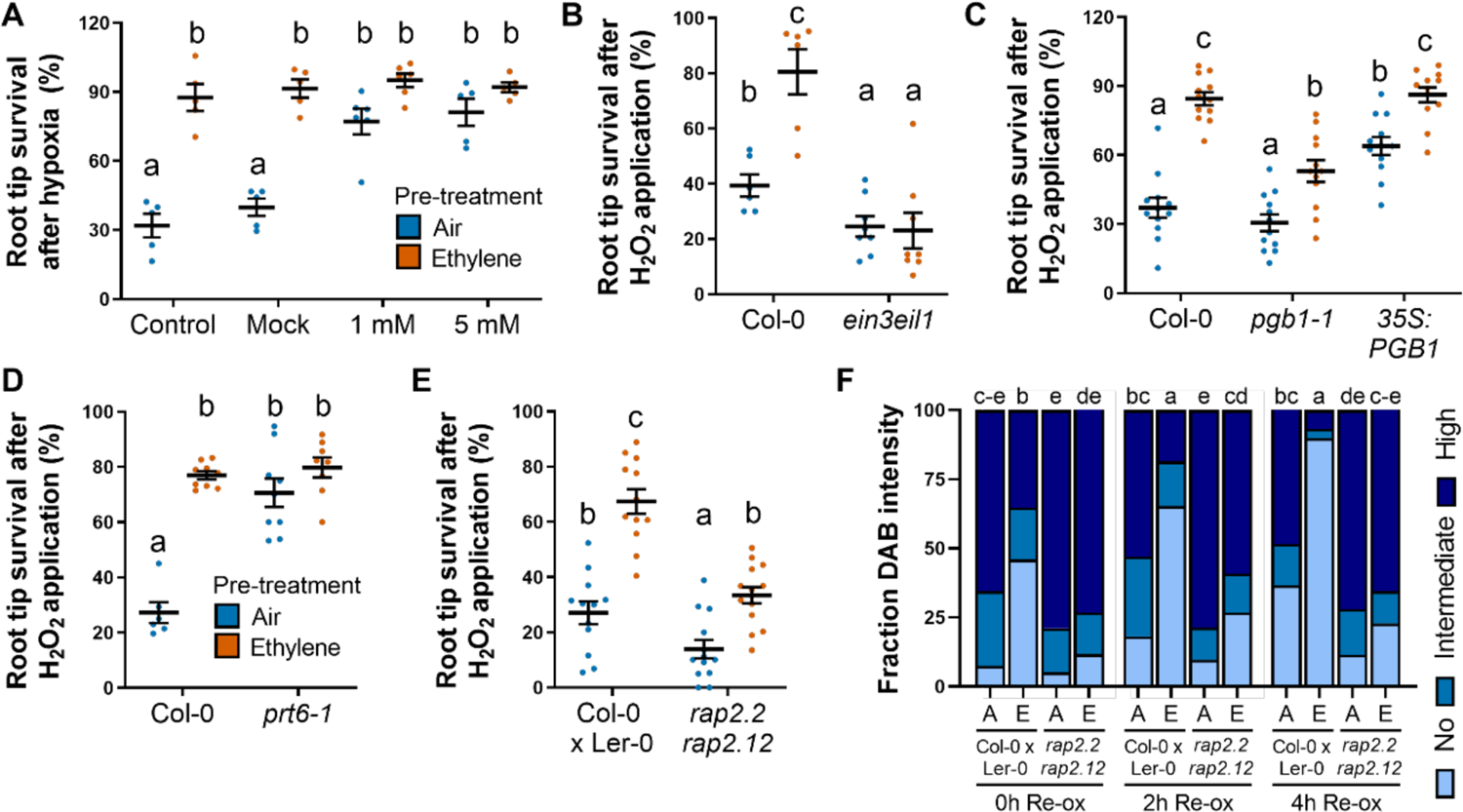
Ethylene improves oxidative stress survival. **(A)** Root tip survival of 4-day old Arabidopsis (Col-0) seedlings after 4 h of air (blue) or ∼5μL·L^-1^ ethylene (orange) treatment in combination with application of mock or 1 mM or 5 mM ROS scavenger potassium iodide (KI), followed by 4 h of hypoxia and 3 days of recovery (n=5 plates containing ∼46 seedlings). (**B-E)** Root tip survival of (B) Col-0 and ethylene insensitive mutant *ein3eil1*, (C) Col-0, *pgb1-1* and *35S:PGB1,* (D) Col-0, N-degron mutant *prt6-1* (Col-0 background) and a (E) Col-0 x Ler-0 cross and ERFVII double mutant *rap2.2rap2.1*2 (Col-0 x Ler-0 background) after 4 h of air (blue) or ∼5 μL·L^-1^ ethylene (orange) treatment followed by application of 5 μL of H_2_O_2_ (6mM) on each single root tip and 3 days of recovery. Statistically similar groups are indicated using the same letter (n=5-12 rows containing ∼23 seedlings). **(F)** Proportion of WT (Col-0 x Ler-0 background) or ERFVII double mutant *rap2.2rap2.1*2 (Col-0 x Ler-0 background) root tips showing H_2_O_2_ (DAB) staining after air (blue) or ∼5 μL·L^-1^ ethylene (orange) treatment followed by 4 h of hypoxia and 2 h of re-oxygenation. Statistically similar groups are indicated per time-point using the same letter (n=146-180 root tips). Letters indicate significantly indistinguishable groups (P < 0.05) based on Tukey’s HSD following a generalized linear model with binomial link function (GLM-binomial). The effect of ethylene on ROS tolerance or DAB staining was tested for with an interaction term in a GLM-binomial with a genotype (wild-type and mutant) and pre-treatment (ethylene-control) effect. Interaction terms for H_2_O_2_ tolerance were *ein3/eil1* (P=7e-5), *pgb1* (P=1e-4), *35S:PGB1* (P=0.002), *prt6-1* (P=3e-8), but not for *rap2.12/2.2* (P=0.21). DAB staining interaction terms were 0 h (P=0.01), 2 h (P=0.04), 4 h (P=9e-6).

We showed previously that ethylene-mediated hypoxia tolerance is controlled by impairment of the PRT6 N-degron pathway through PGB1-mediated NO depletion and subsequent enhanced stability of RAP2.2 and RAP2.12 (Hartman et al., 2019). In the current proteomics data set, PGB1 was the most enriched protein in response to ethylene treatment (Figure 3C). To evaluate whether this mechanism also contributes to ethylene-mediated ROS amelioration, we first assessed root tip survival following H_2_O_2_ application in a *PGB1* knock-down (Hartman et al., 2019) and over-expression line. Ethylene still increased H_2_O_2_ survival in *pgb1-1*, but to a lesser extent than the WT Col-0 (Figure 5C). Moreover, *35S:PGB1* exhibited enhanced oxidative stress survival without an ethylene pre-treatment, suggesting that PGB1 is at least partially involved in ethylene-mediated oxidative stress tolerance. We also examined root tip H_2_O_2_ survival in the *prt6-1* mutant in which proteins with the corresponding N-recognin are stabilized, with a subsequent effect on downstream targets and hypoxia tolerance (Holman et al., 2009; Riber et al., 2015; Zhang et al., 2015).The *prt6-1* root tips showed a strong increase in oxidative stress survival compared to Col-0, such that *prt6-1* seems constitutively ethylene primed (Figure 5D).

NO scavenging by ethylene-induced PGB1 limits ERFVII RAP2.2 and RAP2.12 turnover via the PRT6 branch of the N-degron pathway, aiding hypoxia acclimation (Hartman et al., 2019). We next tested the double null mutant *rap2.2 rap2.12* for ethylene-induced ROS tolerance. Basal levels in H_2_O_2_ tolerance were slightly reduced, but ethylene-enhanced tolerance was unaffected in the double mutant (Figure 5E) relative to the wildtype background. Given that *rap2.2 rap2.12* has reduced hypoxia tolerance (Gasch et al., 2016; Hartman et al., 2019), but did not impact ethylene-mediated oxidative-stress tolerance during normoxia (Figure 5E), we tested whether this line may be altered in ROS homeostasis under actual hypoxia. For this, we used DAB staining to visualize H_2_O_2_ in root tips during hypoxia and reoxygenation (Figure 5F). Indeed, in the wild-type, ethylene increased the number of root tips free of DAB stain and this ethylene benefit was absent in the *rap2.2rap2.12* double mutant. The oxidative stress tolerance assay and ROS profiling under hypoxia suggest that RAP2.2 and RAP2.12 may not directly aid ethylene induced ROS scavenging, but rather that hypoxia acclimation and associated viability mediated by these ERFVIIs (Hartman et al. 2019) is crucial to mount a successful response to reoxygenation. This contrasts with the results for *PGB1* and *PRT6*, suggesting that these proteins, in addition to their role in ERFVII stabilization also directly play a key role in ethylene-mediated ROS scavenging.

### Ethylene rapidly limits root growth which may contribute to enhanced hypoxia tolerance

The transcriptomic analysis revealed both established and novel ethylene-driven gene clusters and genes that are indicative of ethylene-mediated root growth cessation. Indeed, ethylene is a known inhibitor of root growth and development (Le et al., 2001). We hypothesized that energy and resources saved by inhibition of development could, in addition to ROS amelioration, aid in hypoxia tolerance. We first investigated whether the duration of the ethylene pre-treatment used was sufficient to inhibit root growth in our experimental system. A significant reduction in root growth (measured as an increase in primary root length) was observed already after 1 h of ethylene treatment compared to the air control. This root growth cessation was maintained for the entirety of the 4 h ethylene pre-treatment (Supplemental Figure S9).

To separate the effects of growth cessation from other ethylene-mediated tolerance mechanisms we employed the *asa1-1* mutant. ASA1 is a direct target of ERF1 and encodes a rate-limiting step of tryptophan biosynthesis, required for auxin biosynthesis. Ethylene-mediated auxin accumulation via ASA1 leads to inhibition of primary root growth, but this is reported to be absent in *asa1-1* (Mao et al., 2016). The *asa1-1* mutant thus would allow us to apply ethylene and elicit the entire ethylene signaling pathway, but not inhibit root growth. However, in our experimental set-up *asa1-1* had faster root growth rates compared to wild-type, under both control and ethylene conditions. The *asa-1* mutation also did not impact ethylene-mediated root growth inhibition (Figure 6A). Nonetheless, the subsequent increased root growth in *asa1-1*, lead to reduced root tip survival under hypoxia compared to wild type, in both control and ethylene pre-treatment. Furthermore, similar to root growth inhibition, the positive effects of ethylene on survival were maintained in *asa1-1* (Figure 6B). The *asa1-1* mutant, with altered growth rates and altered tolerance, resulted in a continuum of growth-tolerance values with a strong relationship that associated low growth with high survival (Figure 6C).

**Figure 6.**
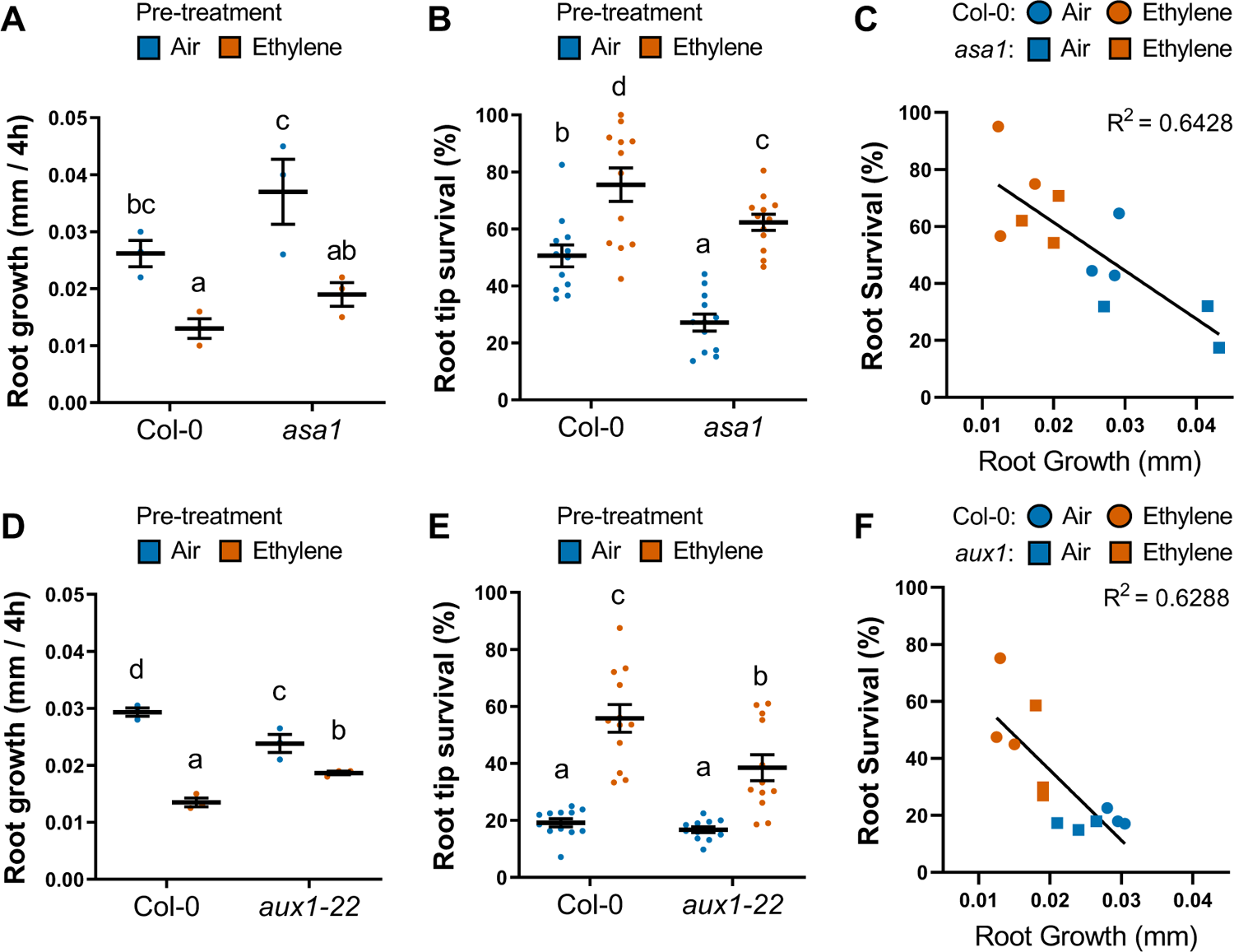
Ethylene-mediated root growth cessation contributes to enhanced hypoxia survival. **(A)** Root growth (increase in root length) of 4-day old Col-0 and *asa1-1* during 4 h of air (blue) or ∼5μL·L^-1^ ethylene (orange) treatment compared to root length at t=0. Data are the median growth rate of single plate of 46 seedlings (N=3). Interaction Pvalue is 0.23. **(B)** Root tip survival of 4-day old Col-0 and *asa1-1* seedlings after 4 h of air (blue) or ∼5μL·L^-1^ ethylene (orange) followed by 4 h of hypoxia and 3 d of recovery. combined data from 3 experiments is shown ((n=12 plates containing ∼46 seedlings). Interaction Pvalue is 0.05. **(C)** Correlation between growth and survival. Mean results of two independent experiments are shown. **(D)** Root growth (increase in root length) of 4-day old Col-0 and *aux1-22* during 4 h of air (blue) or ∼5 μL·L^-1^ ethylene (orange) treatment compared to root length at t=0. Data are the median growth rate of single plate of 46 seedlings (N=3). Interaction Pvalue is 4e--4. **(E)** Root tip survival of 4-day old Col-0 and *aux1-22* seedlings after 4 h of air (blue) or ∼5 μL·L^-1^ ethylene (orange) followed by 3.5 h of hypoxia and 3 days of recovery (n=12 plates containing ∼23 seedlings, combined data from 3 experiments is shown). Interaction Pvalue is 0.01 **(F)** Correlation between growth and survival. Mean results of two independent experiments are shown. Letters indicate significantly indistinguishable groups (P < 0.05) based on Tukey’s HSD following an ANOVA (growth data) or a generalized linear model with binomial link function (GLM-binomial, survival data). Interaction terms were assessed with a GLM-binomial or 2-way ANOVA with a genotype (wild-type and mutant) and pretreatment (ethylene-control) effect.

Manipulation of ASA1 function led to changes in growth independent of ethylene and revealed the relationship between growth and tolerance. To further explore manipulation of ethylene-mediated growth inhibition we utilized the *aux1-22* mutant (Swarup et al., 2007). In the auxin signaling *aux1-22* mutant, ethylene treatment still leads to known ethylene-responsive gene induction, but no longer to auxin-responsive gene induction and subsequent root growth cessation by ethylene in the epidermis (Růžička et al., 2007; Swarup et al., 2007; Vaseva et al., 2018). In our setup ethylene did cause a significant reduction in *aux1-22* root growth, but this inhibition of root growth was considerably less than in Col-0 (Figure 6D). This trend was mirrored in root hypoxia survival. While ethylene still enhanced *aux1-22* hypoxia survival, the effect was significantly reduced relative to the wild-type (Figure 6E). Subsequently, also with *aux1-22,* we found a clear association between low root growth and high survival (Figure 6F). Overall, we demonstrate that growth manipulation, either dependent or independent from ethylene, leads to altered hypoxia tolerance. This suggests that growth cessation contributes to ethylene-mediated tolerance to hypoxia.

## Discussion

In this study we looked for potential genes, proteins and processes that contribute to ethylene-mediated improved hypoxia survival of Arabidopsis root tips. Our results show that ethylene increases both hypoxia and re-oxygenation survival. The data suggest that ethylene may contribute to enhanced tolerance through multiple processes including enhanced hypoxia responses, ROS amelioration responses, and reduction of root growth (Figure 7).

**Figure 7.**
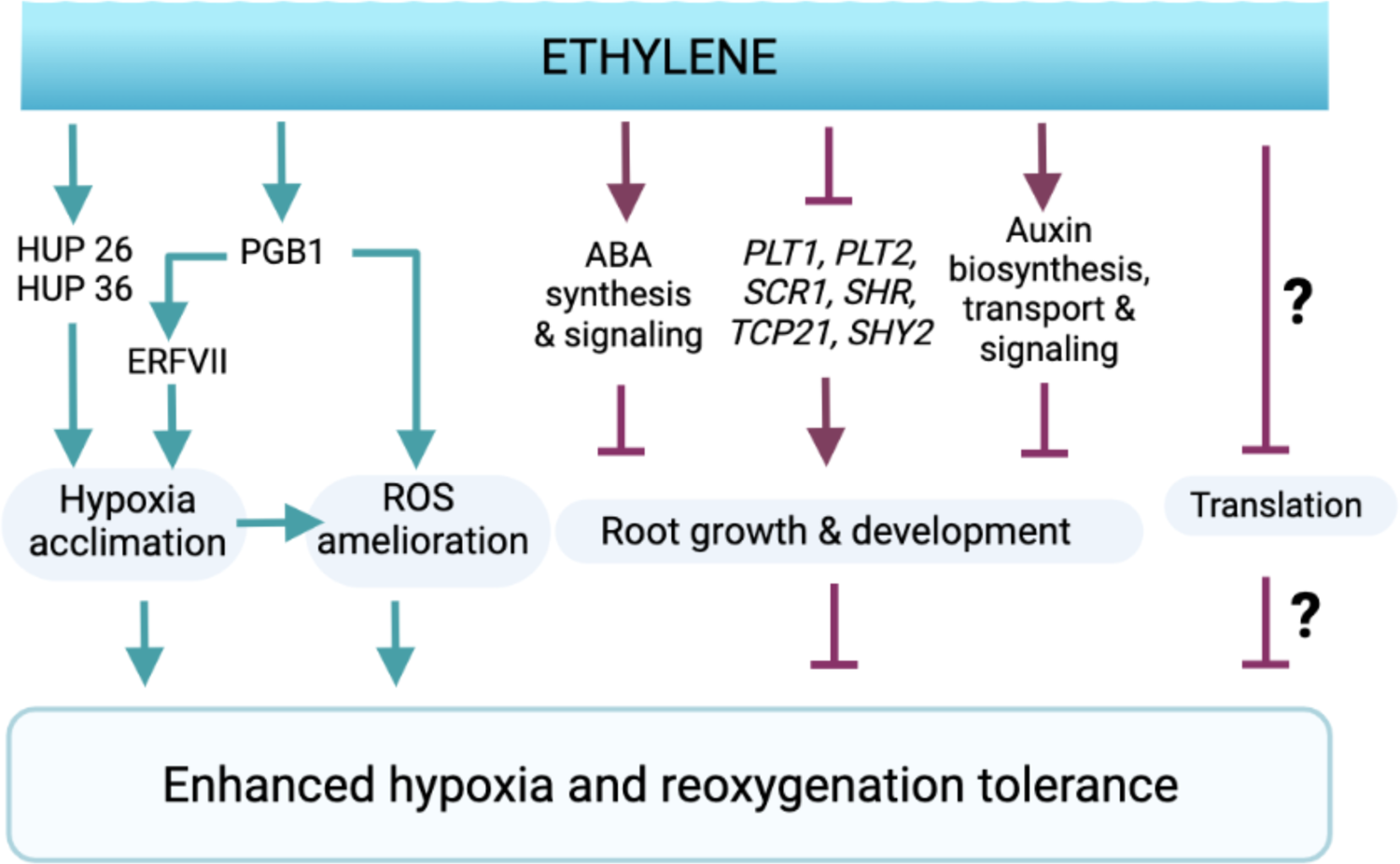
Ethylene modulates a range of acclimation responses to sustain hypoxia and re-oxygenation damage. Proposed model of how ethylene contributes to sustain hypoxia and reoxygenation stress. Ethylene accumulation during flooding prepares cells for impending severe hypoxia via various means: enhanced accumulation of PGB1 (which scavenges NO and stabilizes ERFVIIs) and HUP proteins required for hypoxia acclimation; ROS homeostasis is also mediated by PGB1, either directly via yet unknown mechanisms or indirectly because of ERFVII stabilization and consequent cellular hypoxia protection; Repression of energetically expensive processes of translation and root growth and development. The latter likely involves transcriptional regulation of root developmental regulators and hormone activity. Together the enhanced hypoxia response, ROS amelioration and root growth cessation by ethylene contribute to reduce hypoxia and reoxygenation stress and improve hypoxia survival. Created with BioRender.com

Ethylene has an important role as a distress signal during various environmental stresses. This can be of critical importance for tolerance and has been reported for hypoxia (reviewed in Hartman et al., 2021), salt (Peng et al., 2014), drought, freezing (Wu et al., 2008), and soil compaction stress (Pandey et al., 2021). In our study, the major ethylene-mediated transcriptomic changes occurred during the pre-treatment, without any additional changes upon subsequent hypoxia. This suggests that ethylene-mediated changes occurring before the onset of hypoxia contribute to increased tolerance during hypoxia and re-oxygenation. Indeed, in general, translation and protein synthesis are typically strongly restricted during severe hypoxia stress (Chang et al., 2000; Juntawong et al., 2014). Accordingly, an acclimation period prior to the onset of severe stress can improve subsequent stress tolerance. We propose that for flooding, ethylene could fulfil such a role because inducing protective responses may be energetically challenging to achieve under severe hypoxic stress. Both the transcriptomics and proteomics results suggest that ethylene may also control a general downregulation of translation and growth, which could be adaptive to quickly limit energy expenditure. Ethylene-mediated repression of growth responses and translation could be of great importance to improve survival chances during hypoxia.

Ethylene partially restrains root growth by stimulating auxin synthesis and transport (Růžička et al., 2007; Swarup et al., 2007; Mao et al., 2016). Our results suggest that this ethylene-induced and AUX1-dependent auxin signaling in the root could contribute to subsequent hypoxia survival, as *aux1-22* was more sensitive to hypoxia after ethylene treatment compared to the WT (Figure 6B). While transcriptome profiling identified the classical genes involved in ethylene-mediated root growth, it also identified many growth regulators that were not previously associated with ethylene, such as *UPBEAT1*, *MYB30* and *PLETHORA1/2* (Aida et al., 2004; Tsukagoshi et al., 2010; Mabuchi et al., 2018). Currently, it is unclear whether these are under direct control of ethylene signaling or act indirectly via ethylene-mediated auxin and ROS homeostasis. It also remains to be investigated whether their regulation by ethylene contributes to root growth cessation and hypoxia tolerance. Ethylene-mediated root growth suppression seems universal in the plant kingdom (Visser and Pierik, 2007). That the *asa1* and *aux1-22* root tips, which have altered growth rates either with or without ethylene, have altered hypoxia tolerance or capacity to prime suggests that this growth suppression mediated by early ethylene is a critical strategy for flooding-induced hypoxia survival. The functional consequences of ethylene-mediated growth repression on hypoxia tolerance could therefore be a promising avenue for future research and for the development of flood tolerant crop cultivars.

In addition to root growth cessation and translation, we identified an overrepresentation of genes and proteins related to hypoxia and flooding responses (Figure 2E, Figure 3). The hypoxia response GO term was enriched for ethylene-independent genes as well as for ethylene-enhanced genes, suggesting that there is both an ethylene-dependent and independent hypoxia response in Arabidopsis root tips. Within the ethylene-mediated hypoxia response cluster there were many genes well-known for their role in hypoxia acclimation, such as *PGB1*, *HRE1/2*, *ADH1*, *PDC1* and several *HUPs* (Licausi et al., 2010; Mustroph et al., 2010; Hebelstrup et al., 2012; Mithran et al., 2014; Hartman et al., 2019). Moreover, the proteomics results revealed that ethylene also enhances the protein abundance of PGB1 and HUP36, 26 and 54. In previous studies, constitutive overexpression of single *HUPs* resulted in altered flooding or hypoxia tolerance only in some lines and these effects were variable (Mustroph et al., 2010; Lee et al., 2011). We observed a minor benefit in hypoxia tolerance in *HUP36*, *26* and *54* overexpressors (Supplementary Figure S7). It is likely that a collection of hypoxia responsive genes needs to be activated in concert for achieving substantial beneficial effects for survival. Our results further emphasize that ethylene can promote a subset of hypoxia-responsive genes and proteins already prior to hypoxia that subsequently promote hypoxia survival (Figure 6).

Finally, we identified an overrepresentation of ethylene-enriched genes and proteins related to ROS homeostasis and amelioration (Figure 2E, Figure 3A). Modulation of ROS homeostasis and the induction of ROS signaling has previously been shown to control plant growth and stress responses (Mittler, 2017; Sasidharan et al., 2018). Ethylene-modulation of ROS signaling also controls several flood-adaptive anatomical and growth responses in species such as rice and maize, such as the formation of aerenchyma and adventitious roots (Steffens et al., 2012; Yamauchi et al., 2014; Ni et al., 2019; Qi et al., 2019). In addition, ROS signaling is essential to regulate the transcriptional hypoxia response and mutants showing abnormal ROS signaling are more hypoxia and flood sensitive (Huang et al.; Pucciariello et al., 2012; Gonzali et al., 2015; Yang and Hong, 2015; Sasidharan et al., 2018). Recent reports demonstrate that ROS signaling is partially dependent on ethylene signaling to transcriptionally induce the hypoxia genes *ADH1* and *HRE1* (Hong et al., 2020). However, ethylene-controlled ROS amelioration during flooding, hypoxia and re-oxygenation stress has hardly been investigated (Yeung et al., 2019; Hartman et al., 2021). Prior research has shown that ethylene-insensitive Arabidopsis mutants experience increased damage during re-oxygenation (Tsai et al., 2014). Accordingly, we show that an ethylene pre-treatment improves root tip survival during hypoxia and re-oxygenation stress (Figure 1, Supplemental Figure S1). These results coincide with the observation that hypoxia and re-oxygenation lead to excessive ROS formation in root tips and that ethylene strongly limits this accumulation (Figure 4D-E). Importantly, we demonstrate that enhanced ROS scavenging during and after hypoxia is essential to improve hypoxia survival (Figure 5A), and that ethylene treatment markedly improves the survival of root tips subjected to oxidative stress (Figure 5B-D). Collectively, these data strongly suggest that ethylene facilitates hypoxia and re-oxygenation stress survival through enhanced ROS amelioration (Figure 6).

How ethylene exactly promotes ROS homeostasis to facilitate hypoxia tolerance remains unclear. However, we found that ethylene enhanced transcript levels of genes involved in the Glutathione-AA pathway (Foyer and Noctor, 2011), and changed the abundance of several peroxidases. We also observed that an ethylene treatment alone led to an increase in the total glutathione pool and peroxidase activity which was maintained during subsequent hypoxia (Figure 4A, C). These observations are consistent with reports showing that ethylene promotes peroxidase activity (Gahagan et al., 1968; Mehlhorn, 1990) and glutathione production (Yoshida et al., 2009). Since both glutathione and peroxidases are essential to buffer cellular antioxidant capacity (Gill et al., 2013; Das and Roychoudhury, 2014), it would be interesting to identify whether these changes are required for ethylene-mediated oxidative stress and hypoxia tolerance.

We previously demonstrated that ethylene-induced PGB1 is essential to prevent NO-dependent ERFVII proteolysis and confer subsequent hypoxia tolerance (Hartman et al., 2019). Here we show that PGB1 and the PRT6 pathway are also partially involved in ethylene-mediated ROS tolerance (Figure 5C-E). Notably, enhanced oxidative stress tolerance mediated by ethylene does not require ERF-VIIs (Figure 5E). However, ERFVII loss of function did affect the ethylene mediated ROS amelioration during and after hypoxia (Figure 5F). A similar effect was found in ERFVII single mutants that responded poorly to osmotic stress but were unaffected by hydrogen peroxide treatment (Papdi et al., 2015). We suggest that ERFVIIs are key for hypoxia acclimation, leading to improved viability and therefore indirectly improve the capacity of the cellular machinery for ROS scavenging and homeostasis during reoxygenation stress.

In contrast, PGB1 and PRT6 function are important and likely have a direct effect on ethylene-improved antioxidant capacity. Previous research has linked ethylene, PGB1 and PRT6 to reduced ROS levels during multiple abiotic stresses, including flooding and hypoxia stress. Indeed, overexpression of PGB1 orthologues in maize and soybean led to elevated antioxidant levels, reduced ROS accumulation, enhanced waterlogging and drought stress tolerance and was linked PGB’s NO scavenging capacity (Youssef et al., 2016; Hammond et al., 2020; Mira et al., 2021). In barley, a *prt6* knock out also exhibited improved ROS tolerance and increased performance under abiotic stress (Vicente et al., 2017). Additionally, proteomic analysis of *prt6* Arabidopsis roots showcases elevated expression of three peroxidases and hemoglobin (Zhang et al., 2015), potentially driving the high ROS tolerance observed in *prt6*. Finally, ethylene plays a critical role in ROS amelioration during abiotic stress responses (Wu et al., 2008; Peng et al., 2014; Tsai et al., 2014), and antioxidant activity is a crucial driver for enhanced abiotic stress tolerance (Arbona et al., 2008; Fukao et al., 2011; Yeung et al., 2018). Collectively, these results suggest that ethylene-mediated ROS amelioration could be an important general trait for abiotic stress resilient crop breeding and identifies PGBs and PRT6 as potential regulatory targets.

Acute hypoxia as a result of flooding places a heavy burden on plant cells because energy and carbohydrate generation are strongly impaired while these are required to mount adaptive responses and sustain the accumulation of highly damaging ROS. We show the early flooding signal ethylene improves hypoxia survival by regulating a range of processes that facilitate hypoxia acclimation and prevent post-hypoxic injury. These ethylene-controlled processes modulated at the transcriptional and protein level include hypoxia responses, ROS metabolism, growth cessation, and possibly the downregulation of translation and leads to subsequent enhanced ROS amelioration and hypoxia survival (Figure 7). Finally, we show that ethylene improves oxidative stress survival and propose that this process is pivotal for how ethylene confers hypoxia tolerance in plants.

## Supporting information

SupplementalData_Liu_others

## Acknowledgements

We thank Malcolm Bennett and Bipin Pandey for providing seeds of *aux1-22* (Swarup et al., 2007), Angelika Mustroph for *rap2.2 rap2.12* and its wildtype background crosses (Gasch et al., 2016), Thomas Nietzel and Markus Schwarzländer for *mt-roGFP2-Orp1* (Nietzel et al., 2019). We acknowledge Dr. Michael J. Deery of Cambridge Centre for Proteomics, Cambridge University, for assistance with mass spectrometry, Nienke van Dongen, Emilie Reinen and Ankie Ammerlaan for technical support.

## Funding

This work was supported by the Netherlands Organization for Scientific Research; 831.15.001 and 019.201EN.004 to S.H., 824.14.007 to L.A.C.J.V, S.M., BB.00534.1 to R.S and ALWOP.419 to HvV and RS. ZL was supported by a China Scholarship Council grant No. 201406300100. Research at Rothamsted was funded by CSIA grant 18-6 awarded to HZ and the BBSRC Tailoring Plant Metabolism Institute Strategic Grant BBS/E/C/000I0420. MH and JBS were supported by US National Science Foundation (MCB-1716913).

## Author contributions

ZL, SH, HvV, HZ designed and performed research and analyzed the data. HACFL, SM, FB, FD, PS, MH, TR, KLH performed research and analyzed the data. JBS, FLT, LACJV and RS designed research. LACJV and RS coordinated the research and wrote the manuscript together with ZL, SH and HvV. All authors saw and commented on the manuscript.

## Materials and Methods

### Plant material and growth conditions

*Arabidopsis thaliana* wild type lines Col-0, Col-0 x Ler-0 and null mutants *ein3-1eil1-1*, *PRT6* defect *prt6-1* (SAIL 1278_H11), *PGB1* knock-down *pgb1-1* (SALK_058388) and over-expression line *35S::PGB1*, ERFVII double mutant *rap2.2rap2.1*2 (Col-0 x Ler-0 background) were previously described (Hartman et al., 2019), and sterilized for 3 h in a mixture of bleach (30 mL) and concentrated hydrochloric acid (1.5 mL, Sigma-Aldrich). The *pgb1-1* mutant lacks ethylene-induced *PGB1* expression and lower hypoxia induced *PGB1* expression (relative to wild-type) (Hartman et al., 2019). Seeds were sown on sterile, square petri dishes (120×120×17mm, Greiner Bio One) on 1% plant agar (25 mL, Duchefa Biochemie B.V., P1001) supplemented with ¼ Murashige & Skoog (MS) (Duchefa Biochemie B.V., M0245). Forty-six seeds were sown in two rows per plate and plates were sealed with Leucopore tape (12.5 mm, Duchefa Biochemie B.V., L3302). After 4 days of stratification at 4°C in the dark, plates were placed vertically in climate-controlled growth chambers with a short day photoperiod (daytime: 9-17 h, light intensity: ∼120 μmol·m^-2^·s^-1^, humidity: 70%, temperature: 21°C daytime and 18°C nighttime). For recovery, plates were kept in the dark after hypoxia treatment and then moved back to the same short day growth conditions when the light was switched off in the climate chamber at the end of the day.

### Ethylene and hypoxia treatment conditions

For ethylene treatments, seedlings grown on square plates with lids removed were placed vertically inside closed glass desiccators injected with specific ethylene concentrations (light intensity within desiccators: ∼5 μmol·m^-2^·s^-1^, room temperature). Seedlings placed in glass desiccators without ethylene were used as control. Commercially purchased gaseous ethylene was diluted to desired concentrations and injected into glass desiccators. After 30 minutes, gas samples were taken from the desiccators to confirm the ethylene concentration by gas chromatography (Syntech Spectras GC955). For hypoxia treatments, seedlings grown on square plates with lids removed were placed vertically inside closed glass desiccators, which was flushed with gaseous nitrogen at a rate of 2 L·min^-1^ in the dark (typically reaching 0.00% O_2_ after 40-50 min of treatment, see Hartman et al., 2019). Seedlings placed in desiccators in which air was flushed at a similar rate were used as control.

### H*_2_*O*_2_* treatments

H_2_O_2_ (30% w/w, Merck KGaA) was diluted into 1/4 MS to achieve desired concentrations and 5 μL of H_2_O_2_ solution was applied to each root tip in the dark. Plates were kept horizontal for 15 minutes to allow the solution to be absorbed by the root tips.

### Evans blue staining for cell viability and visualization

For each treatment per replicate, around 20 seedlings were randomly taken from the plates for Evans blue staining of cell integrity at desired time points during hypoxia and subsequent re-oxygenation. Seedlings were incubated in 0.25% aqueous Evans blue solution for 15 minutes in the dark (room temperature), then washed three times with Milli-Q (MQ) water to remove excess dye. Olympus BX50WI was used for visualization and images were taken under 10X objective lens.

### Root tip survival and growth quantification and data analysis

Primary root tip survival was scored according to root tip re-growth after 3 days of recovery following hypoxia treatment (Hartman et al., 2019). Survival rate was calculated as the percentage of seedlings that showed root tip re-growth out of the 23 seedlings per row grown vertically on an agar plate. Root tip growth was measured using ImageJ software. Statistical significance was analyzed as indicated in figure legends.

### Micro-array sample preparation and analysis

Samples were harvested at desired time-points (Figure 2A) and snap frozen in liquid N_2_ immediately. Total RNA was isolated using the RNeasy mini kit (Qiagen, Germany). Micro-array was performed commercially by Macrogen (Seoul, South Korea). RNA purity and integrity were evaluated by ND-1000 Spectrophotometer (NanoDrop, Wilmington, USA), Agilent 2100 Bioanalyzer (Agilent Technologies, Palo Alto, USA). RNA labeling and hybridization were performed by using the Agilent One-Color Microarray-Based Gene Expression Analysis protocol (Agilent Technology, V 6.5, 2010). Briefly, RNA was linearly amplified and labeled with Cy3-dCTP, purified using the RNeasy mini Kit, and then measured using NanoDrop ND-1000. 1650ng of labeled cRNA was fragmented by adding 11 μL 10x blocking agent and 2.2 μL of 25x fragmentation buffer and subsequently heated to 60°C for 30 minutes. Finally, 55μL 2x GE hybridization buffer was added to dilute the labeled cRNA. 100μL of hybridization solution was dispensed into the gasket slide and assembled to the Agilent SurePrint HD Arabidopsis GE 4X44K Microarrays (Agilent®). Slides were incubated for 17 h at 65°C in an Agilent hybridization oven, then washed at room temperature using the Agilent One-Color Microarray-Based Gene Expression Analysis protocol (Agilent Technology, V 6.5, 2010). The hybridized array was immediately scanned with an Agilent Microarray Scanner D (Agilent Technologies, Inc.) Results were extracted using Agilent Feature Extraction software v11.0 (Agilent Technologies) and can be found in ArrayExpress under accession number E-MTAB-11231 (https://www.ebi.ac.uk/arrayexpress/experiments/E-MTAB-11231).

### Gene expression analysis

For by gene expression estimates the Arabidopsis GE 4X44K was first matched to the most recent Arabidopsis annotation, Araport11, by the BLAST algorithm. The results were analyzed using the limma R package. Here the background was correct with a moving minimum of 3×3 grids of spots and the expression was quantile normalized. Differentially expressed genes (Padj. < 0.001) were placed in similarly regulated groups by calculating pairwise Euclidean distances followed by agglomerative hierarchical clustering using wards squared method. Gene ontology enrichment was determined with the GOstats R library.

### Proteomics sample preparation and analysis

Plant growth material and conditions: Seedlings grown for proteomics harvests (*Arabidopsis thaliana* seeds of ecotype Col-0) were sown in 3 rows at high density (∼15-20 seeds/cm) on sterile square petri dishes containing 40 ml autoclaved and solidified ¼ MS, 1% plant agar without additional sucrose, using a pipette after wet surface seed sterilization (incubation in 50% EtOH, 5% Bleach for 10 minutes, followed by 7 washing rounds with Autoclaved MQ water). The plates were left to dry for 30 minutes after sowing. Petri dishes were sealed with gas-permeable tape (Leukopor, Duchefa) and stratified at 4°C in the dark for 4 days. Seedlings were grown vertically on the agar plates under short day conditions (9:00 – 17:00, T= 20°C, Photon Flux Density = ∼120 μmol m^-2^s^-1^, RH= 70%) for 5 days. Samples were harvested over 5 independent experiments; 10 plates were pooled per biological replicate. Air and ethylene treatments were performed as described above.

Sample preparation, TMT labeling and data analysis: Protein extraction, quantification, reduction and alkalization was done as described previously (Zhang et al., 2015). Protein precipitation was done by the methanol/chloroform method described by (Zhang et al., 2018). Sequential trypsin digestion was done according to (Zhang et al., 2015). Peptide concentration was determined using a PierceTM Quantitative Colorimetric Peptide Assay kit (23275, Thermo Scientific). 100µg peptide aliquots were labelled using TMT10plex^TM^ (90110, Thermo Scientific). 5 biological replicates were performed: Air-treated samples were labelled with TMT10 −126, −127N, −127C, −128N, −128C and ethylene-treated samples were labelled with −129 N, −129C, −130N, −130C, −131. Peptides were mixed equally and an aliquot corresponding to half of peptides was separated by high pH reverse-phase chromatography using a Waters reverse-phase Nano column as described in (Zhang et al., 2015). All LC-MS/MS experiments were performed as previously (Zhang et al. 2018) using a Dionex Ultimate 3000 RSLC nanoUPLC (Thermo Fisher Scientific Inc, Waltham, MA, USA) system and a Orbitrap Fusion Lumos Tribrid Mass Spectrometer (Thermo Fisher Scientific Inc, Waltham, MA, USA).

Raw data were searched against the TAIR10 database using MASCOT v.2.4 (Matrix Science, London, UK) and PROTEOME DISCOVERER^TM^ v.1.4.1.14 as described previously (Zhang et al., 2015), employing Top 10 peaks filter node and percolator nodes and reporter ions quantifier with Trypsin enzyme specificity with a maximum of one missed cleavage. Carbamidomethylation (+57.021 Da) of cysteine and TMT isobaric labelling (+229.162 Da) of lysine and N-termini were set as static modifications while the methionine oxidation (+15.996) were considered dynamic. Mass tolerances were set to 20 ppm for MS and 0.6 Da for MS/MS. For quantification, TMT 10plex method was used and integration tolerance was set to 2mmu, Integration Method was set as centroid sum. Purity correction factor were set according the TMT10plex (90110) product sheet (Lot number: SA239883). Each reporting ion was divided by the sum of total ions and normalized by medians of each sample. Log transformation ensured a homogeneity and normal distribution of the variances. Statistical differences between air and ethylene samples were obtained using two-sample t-test of log_2_ transformed data, considering variation of quantification as a weighting factor. Only proteins represented by two or more peptides were considered for further analysis. All proteomics data is available in the Supplemental data (sheet 8-10). MS proteomics data have been deposited to the ProteomeXchange Consortium via the PRIDE partner repository (Vizcaíno et al., 2016). R language was used for Ontology (GO) enrichment analysis.

### Antioxidant quantification

Glutathione and Ascorbic acid content were quantified using the GSH-GLO Glutathione Assay Kit (Promega, Madison, USA) and Megazyme kit (K-ASCO 04/19, Wicklow, Ireland), respectively. Briefly, ∼600 root tips were harvested per sample through flash freezing after treatment. The manufacturer’s protocols were adapted for extraction and quantification as previously described (Yeung et al., 2018). Glutathione and ascorbic acid values were normalized by total soluble protein content per sample. Protein content was measured using the Pierce BCA Protein Assay Kit (Thermo-Fisher).

Peroxidase activity was quantified using the Peroxidase Activity Assay Kit (Sigma-Aldrich). Briefly, ∼600 root tips were harvested per sample and rapidly homogenized with 100 mL of Assay Buffer. Sample supernatant (50 mL) was incubated with Peroxidase Master Reaction Mix (50 mL) in 96-well plate and incubated at 37°C for 10 minutes. Colorimetric measurements (absorbance at 570 nm) were performed every 3 minutes for a total time of 18 minutes and a standard curve (0 to 10 nmol H_2_O_2_) was included. Peroxidase activity was calculated by dividing the amount (nmol) of H_2_O_2_ reduced between T_initial_ and T_final_ by the measuring time (18 min) and sample volume (100 mL).

### H*_2_*O*_2_* visualization

3,3’-Diaminobenzidine (DAB) staining: DAB is oxidized by H_2_O_2_ in the presence of peroxidases to generate a dark brown precipitate (Thordal-Christensen et al., 1997). Seedlings were incubated with 1 mg·mL^-1^ DAB (Sigma-Aldrich) in 20 mM 2-ethanesulfonic acid (MES, Sigma-Aldrich) buffer (pH 6.2) supplemented by 10 units·mL^-1^ peroxidase from horseradish (Sigma-Aldrich) for 1 h. Seedlings were then rinsed with MES buffer for 1 minute twice before imaging (Dubreuil-Maurizi et al., 2011). Olympus BX50WI was used to visualize DAB staining and images were taken under 10X objective lens.

Confocal imaging of mt-roGFP2-Orp1: to visualize subcellular levels of H_2_O_2,_ confocal imaging of 5-day old *mt-roGFP2-Orp1* Arabidopsis seedlings was performed right after experimental time-points with a Zeiss Observer Z1 LSM700 confocal microscope (oil immersion, 40x objective Plan-Neofluar N.A. 1.30). RoGFP2-Orp1 was excited sequentially at 405 and 488 nm and emission was recorded at 505–535 nm. The 405/488 ratio within root tips was determined in similar areas of ∼5000 μm^2^ using ICY software (http://icy.bioimageanalysis.org).

---

**Supplemental Table 1.** The effect of ethylene on MC-protein abundance. Table showing the normalized ratios of N-terminal MC-initiating proteins detected in the proteomics dataset in seedling root tips by ethylene, expressed as the protein fold change ratio of ethylene treated seedlings compared to air (**Bold** indicates p<0.05, Student’s t-Test, n=5).

| Gene | Description | Ratio<br>ethylene vs air | p value |
| --- | --- | --- | --- |
| AT4G14430.1 | Indole-3-butyric acid response 10 | 1.107 | 0.060 |
| AT3G51640.1 | Stress response NST1-like protein | 1.107 | 0.730 |
| AT4G24020.1 | NIN like protein 7 | 1.088 | 0.329 |
| AT2G26110.1 | Protein of unknown function (DUF761) | 1.070 | 0.668 |
| AT5G54830.1 | DOMON domain-containing protein / dopamine<br>beta-monooxygenase N-terminal domain | 1.065 | 0.274 |
| AT5G65010.2 | Asparagine synthetase 2 | 1.049 | 0.126 |
| AT3G24090.1 | Glutamine-fructose-6-phosphate transaminase;<br>sugar binding; transaminases | 1.041 | 0.552 |
| AT2G33230.1 | Yucca 7 | 1.037 | 0.574 |
| AT4G28820.1 | HIT-type Zinc finger family protein | 1.030 | 0.459 |
| AT1G06570.1 | Phytoene desaturation 1 | 1.012 | 0.894 |
| AT3G47340.1 | Glutamine-dependent asparagine synthase 1 | 1.002 | 0.991 |
| AT4G16845.1 | VEFS-Box of polycomb protein | 0.999 | 0.995 |
| AT5G10240.1 | Asparagine synthetase 3 | 0.984 | 0.555 |
| AT1G65520.1 | Delta(3), delta(2)-enoyl coa isomerase 1 | 0.977 | 0.684 |
| AT1G22950.1 | 2-oxoglutarate (2OG) and Fe(II)-dependent<br>oxygenase superfamily protein | 0.977 | 0.582 |
| AT2G41900.1 | CCCH-type zinc finger protein with ARM repeat<br>domain | 0.973 | 0.561 |
| AT4G14440.1 | 3-hydroxyacyl-coa dehydratase 1 | 0.972 | 0.891 |
| AT3G03773.2 | HSP20-like chaperones superfamily protein | 0.959 | 0.524 |
| AT3G12600.1 | Nudix hydrolase homolog 16 | 0.957 | 0.419 |
| AT5G48180.1 | Nitrile specifier protein 5 | 0.946 | 0.383 |
| AT2G03667.1 | Asparagine synthase family protein | 0.946 | 0.276 |
| AT4G09320.1 | Nucleoside diphosphate kinase family protein | 0.925 | 0.219 |
| AT1G63610.2 | Unknown protein | 0.923 | 0.144 |
| AT3G63220.2 | Galactose oxidase/kelch repeat superfamily protein | 0.907 | 0.166 |
| AT4G02860.1 | Phenazine biosynthesis phzc/phzf protein | 0.880 | 0.160 |
| AT2G07640.1 | NAD(P)-binding Rossmann-fold superfamily<br>protein | 0.836 | 0.134 |
| AT5G12850.1 | **CCCH-type zinc finger protein with ARM repeat**<br>**domain** | 0.816 | 0.000 |
| AT2G10450.1 | **14-3-3 family protein** | 0.805 | 0.034 |
| AT2G26470.1 | **Embryonic stem cell-specific 5-**<br>**hydroxymethylcytosine-binding protein** | 0.785 | 0.030 |

**Supplemental Table 2.** Comparison of identified proteins regulated by the PRT6 N-degron pathway and ethylene. The table shows the normalized ratios of protein abundance significantly enriched in both N-degron pathway mutants *prt6-1* and *ate1ate2* (adapted from (Zhang et al., 2015)), compared to the protein fold change of those treated with ethylene compared to air (**Bold** indicates p<0.05, Student’s t Test). *Indicates whether the genes encoding for these proteins are direct targets of ethylene transcriptional regulator EIN3 (Chang et al., 2013).

| Gene | Description | Ratio<br>*prt6*:<br>Col-0 | Ratio<br>*ate1/2*:<br>Col-0 | Ratio<br>ethylene : air<br>(in Col-0) | EIN3<br>target |
| --- | --- | --- | --- | --- | --- |
| AT4G27450.1 | **HUP54, Aluminium induced protein with YGL and LRDR motifs** | 3.826 | 6.642 | **1.209** | Yes* |
| AT2G16060.1 | **PGB1, Phytoglobin 1** | 3.218 | 5.200 | **6.403** | No |
| AT3G03270.1 | HRU1, Hypoxia-Responsive Universal Stress Protein 1 | 2.143 | 3.373 | 1.004 | Yes* |
| AT5G19550.1 | ASP2, Aspartate aminotransferase 2 | 1.364 | 1.629 | 0.954 | No |
| AT5G26280.1 | TRAF-like family protein | 1.344 | 1.490 | 1.085 | No |
| AT1G52070.1 | Mannose-binding lectin superfamily protein | 1.314 | 1.201 | 0.891 | No |
| AT5G63550.2 | **DEK domain-containing chromatin associated protein** | 1.279 | 1.404 | **1.256** | No |
| AT3G12580.1 | HSP70, heat shock protein 70 | 1.198 | 1.319 | 1.001 | No |
| AT5G61210.1 | SNP33, soluble N-ethylmaleimide-sensitive factor adaptor protein 33 | 1.194 | 1.331 | 1.103 | No |
| AT5G12110.1 | Glutathione S-transferase, C-terminal-like; Translation elongation factor | 1.152 | 1.454 | 1.191 | No |
| AT4G08770.1 | PRX37, Peroxidase superfamily protein | 1.127 | 1.349 | 1.120 | No |
| AT3G07720.1 | Galactose oxidase/kelch repeat superfamily protein | 1.126 | 1.379 | 1.072 | No |

## Supplemental Data

**Supplemental Figure 1.**
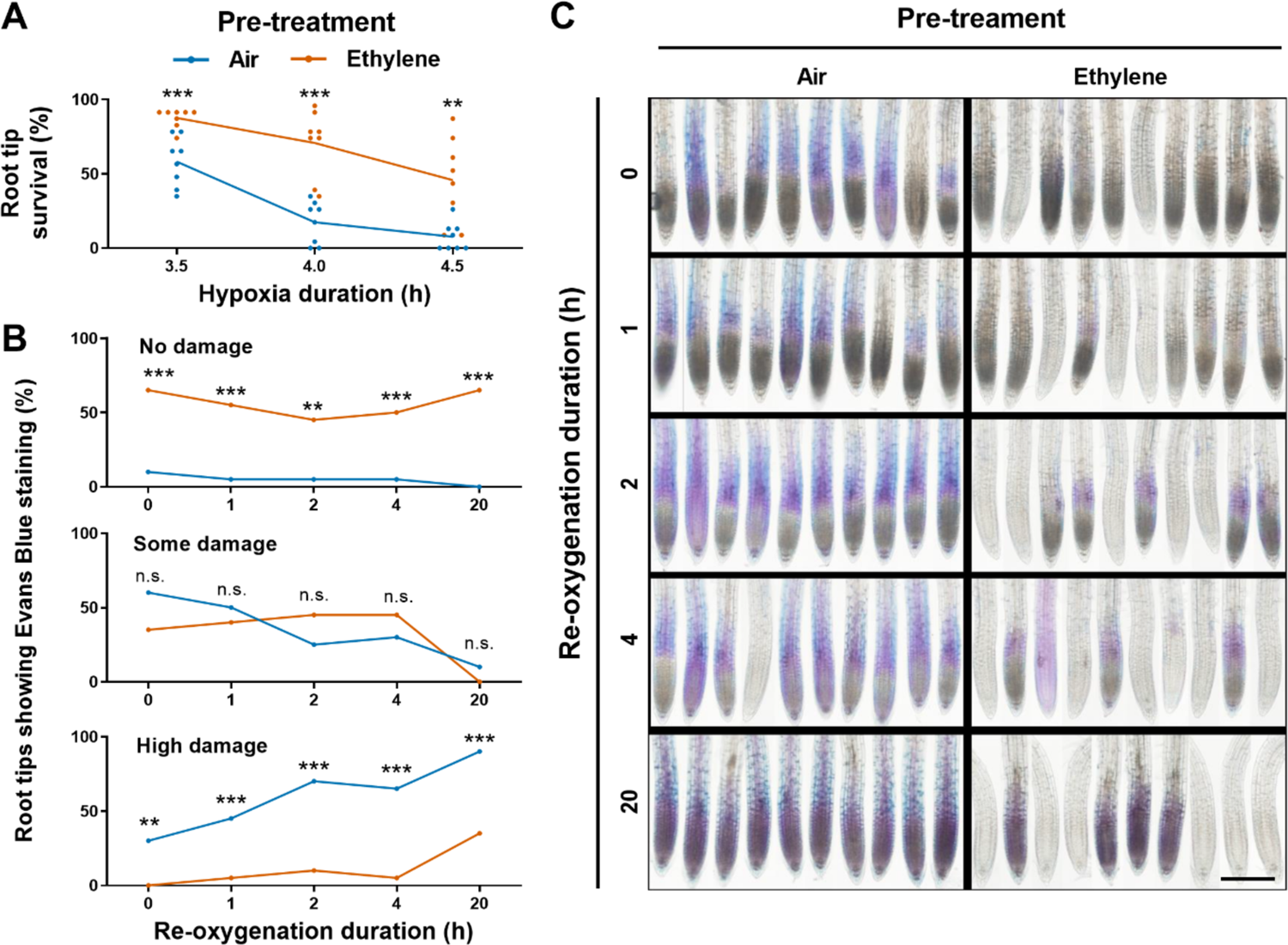
Ethylene pre-treatment improves cell viability during hypoxia and re-oxygenation. **(A)** Root tip survival of 5-day old Arabidopsis (Col-0) seedlings after 4h of air (blue) or ∼5 μL·L^-1^ ethylene (orange) treatment followed by hypoxia and 3 days of recovery (n=8 rows of 23 seedlings). **(B)** Classification and **(C)** visualization of Evans Blue (EB) staining for impaired cell membrane integrity in 4-day old seedling root tips after 4 h of pre-treatment with air or ∼5μL·L^-1^ ethylene followed by 4.5 h hypoxia and re-oxygenation time points. Classification in (B), no damage = no EB staining, some damage = detectable EB staining, high damage = clear root-wide EB staining in elongation zone and root apical meristem, (C) scale bar = 100 µm. Asterisks indicate significant differences between air and ethylene per time point (n.s. - not significant, **p<0.01, ***p<0.001, generalized linear model with a binomial error structure, n=20 root tips in B, n=10 random samples in C).

**Supplemental Figure 2.**
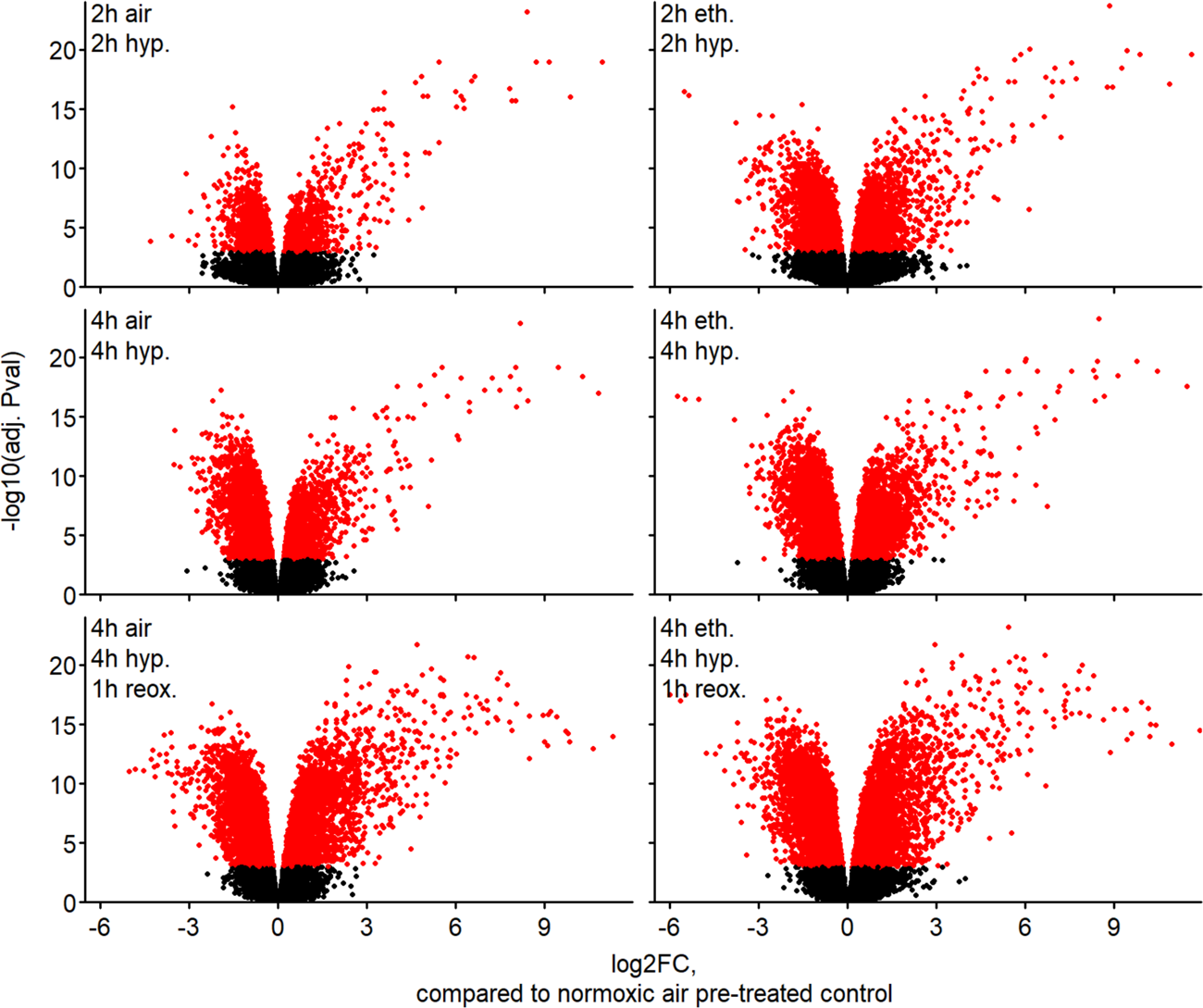
Volcano plots of hypoxia and re-oxygenation transcriptome responses of ethylene and air pre-treated seedlings. Fold changes are compared to the normoxic treated control seedlings of the corresponding timepoint. The genes indicated in red are considered to be differentially regulated (P < 0.001).

**Supplemental Figure 3.**
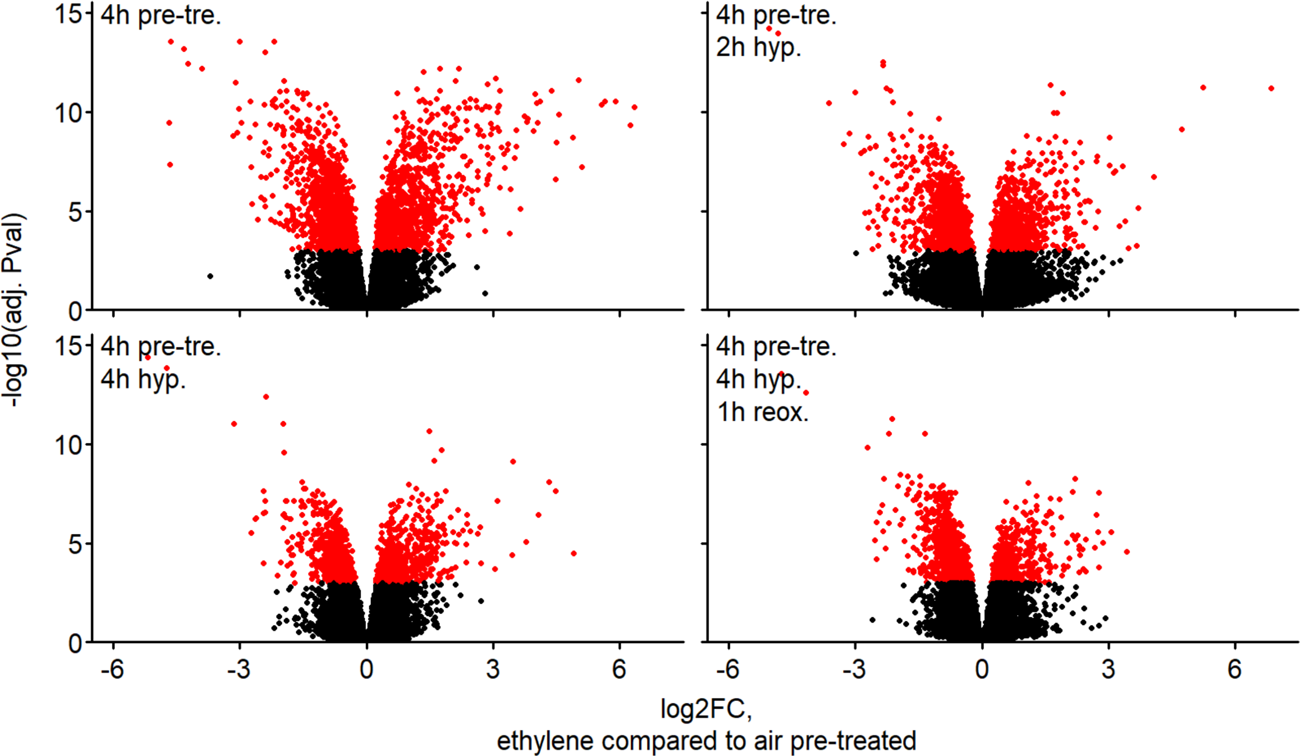
Volcano plots showing the difference between transcriptomes of ethylene and air pre-treated seedlings over the course of the pre-treatment, hypoxia and re-oxygenation. Fold changes represent direct comparisons between ethylene and air pre-treated plants within each timepoint. The genes in red are considered differentially regulated (P < 0.001)

**Supplemental Figure 4.**
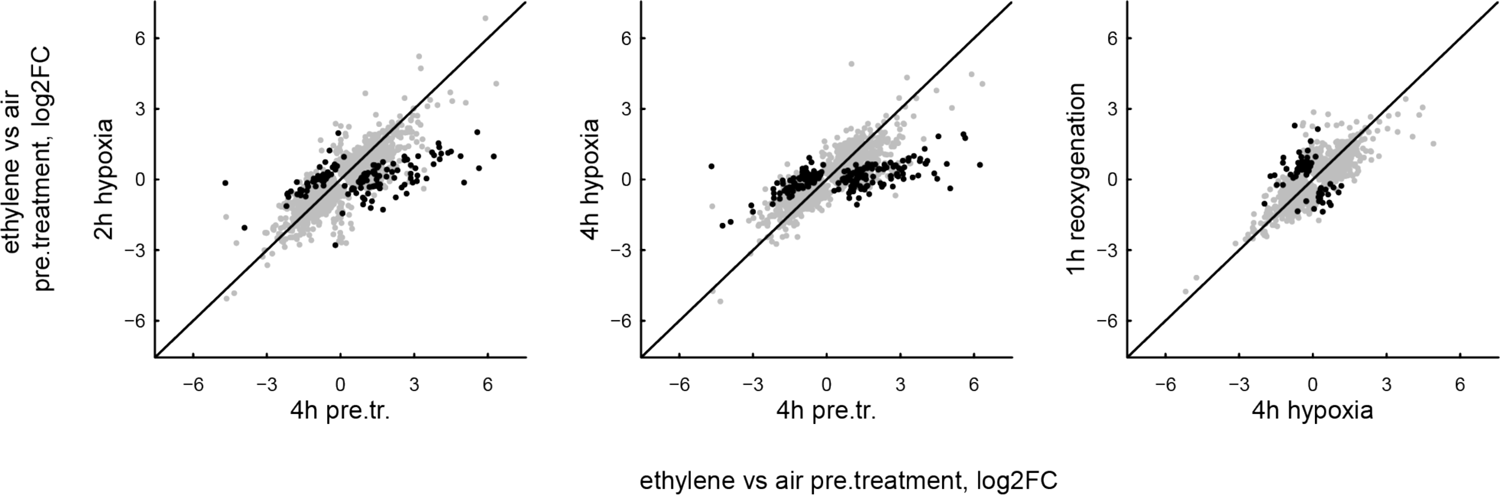
Ethylene specific transcriptional responses during hypoxia and re-oxygenation. Scatter plots showing the log_2_FCs of the ethylene pre-treatment effect for the hypoxia or re-oxygenation timepoints on the y-axis with the FCs of the preceding condition, pre-treatment or hypoxia respectively, on the x-axis. Genes that are differentially expressed in at least one timepoint are shown. Black dots represent genes responded significantly different to the pre-treatment at the compared timepoints (P < 0.001).

**Supplemental Figure 5.**
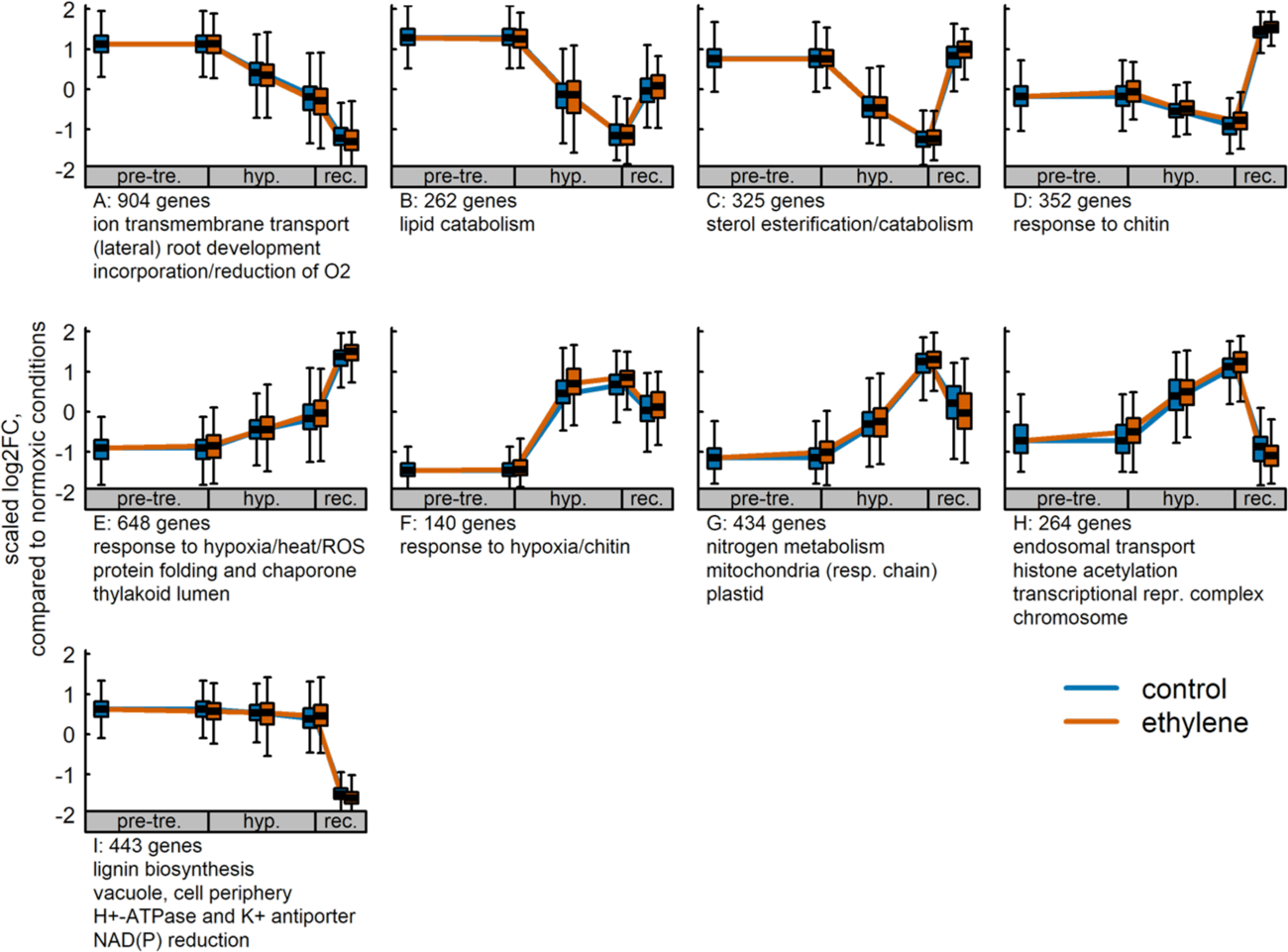
Transcriptional changes that are independent of ethylene pre-treatment. Clusters from hierarchical clustering of genes that respond to hypoxia (p <0.001) but show no differences based on pre-treatment (P > 0.05). Key processes and functions enriched in each cluster based on gene ontology (GO) enrichment. Full lists of GO terms are available in the Supplemental Data (Sheet 5-7). Shown FCs are relative to the normoxic control of the corresponding timepoint, and were mean and standard deviation scaled by gene.

**Supplemental Figure 6.**
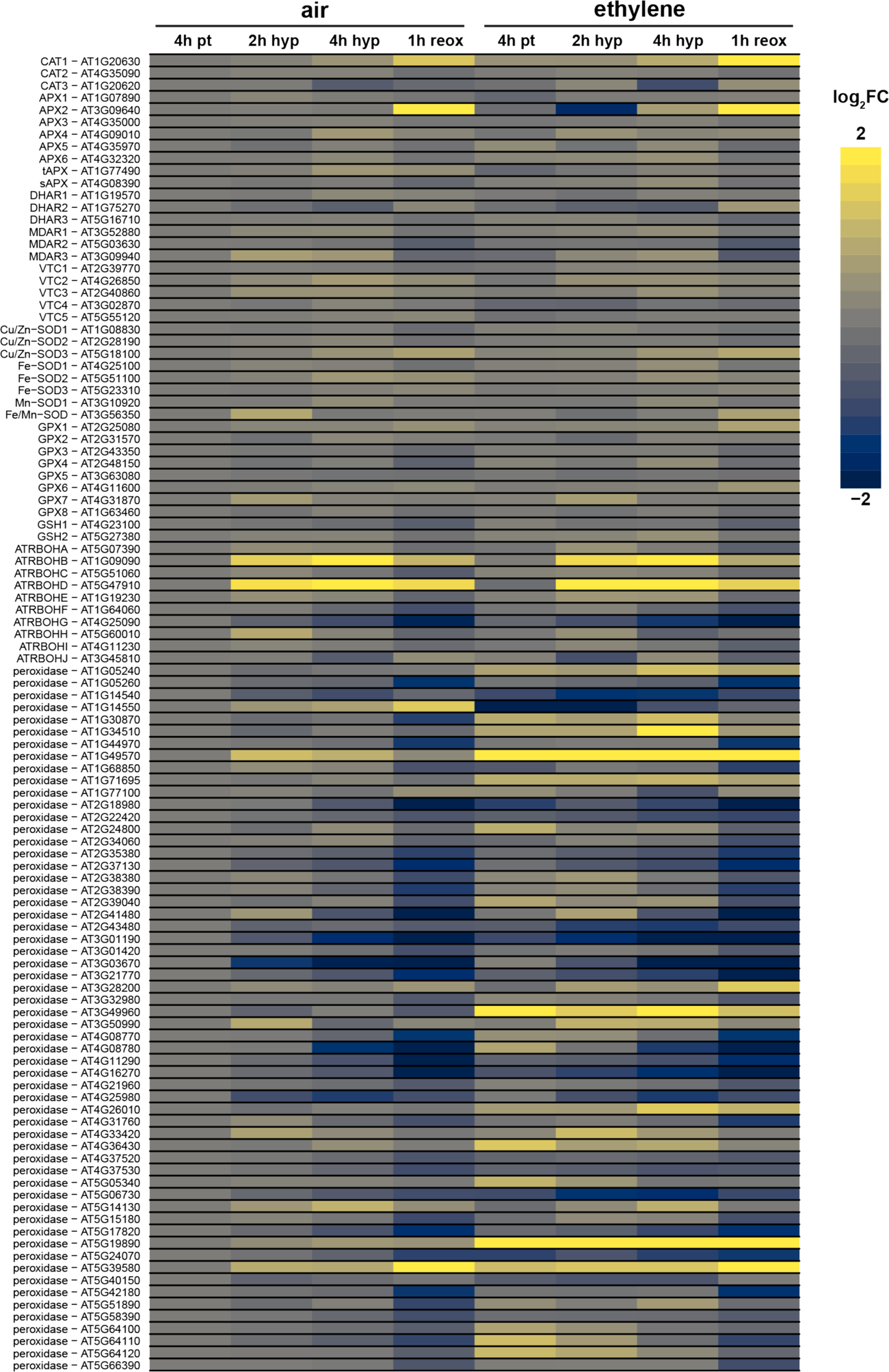
Several key ROS homeostasis gene-families respond differently to ethylene treatment. Heatmap showing the Log_2_FCs compared to the normoxic control of the corresponding time point. Key ROS homeostasis gene-families are shown, including catalase, ascorbate peroxidases and reductase, ascorbate biosynthesis, superoxide disumutases, glutathione peroxidases, glutathione biosynthesis, NADPH-oxidases (RBOHs) and the peroxidases.

**Supplemental Figure 7.**
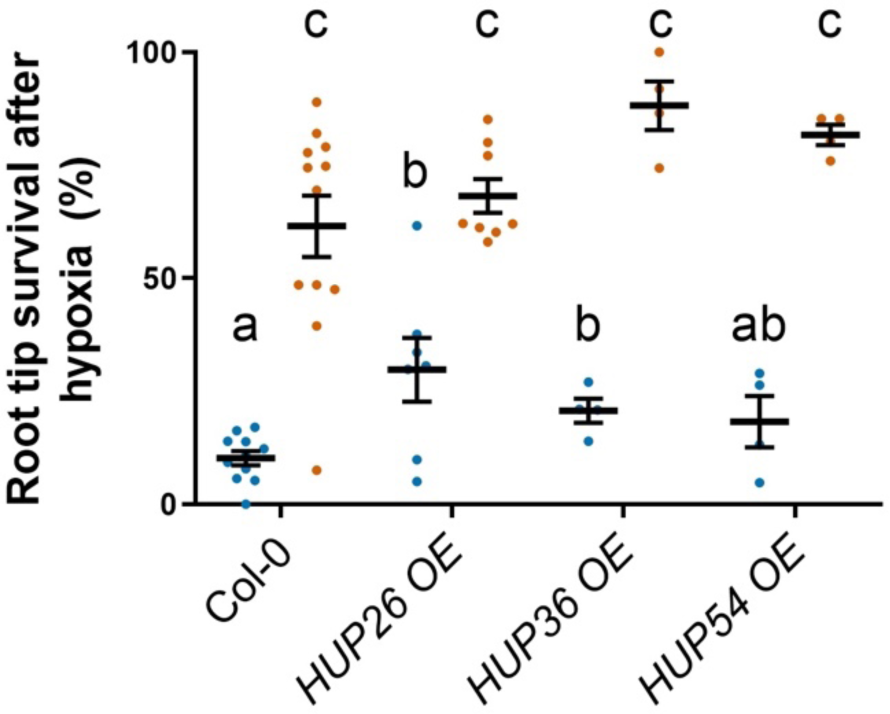
Ethylene-mediated increase in HUPs might contribute to hypoxia survival of Arabidopsis root tips. Seedling root tip survival of Col-0 and 3 over expression lines of *HUP26*, *HUP36* and *HUP54* after 4 h of pre-treatment with air (blue dots) or ethylene (orange dots) followed by 3.5 h of hypoxia and 4 days of recovery. Values are relative to control (normoxia) plants. Statistically similar groups are indicated using the same letter (Error bars are SEM, p<0.05, 2-way ANOVA, Tukey’s HSD, n=4-12 rows of ∼23 seedlings.

**Supplemental Figure 8.**
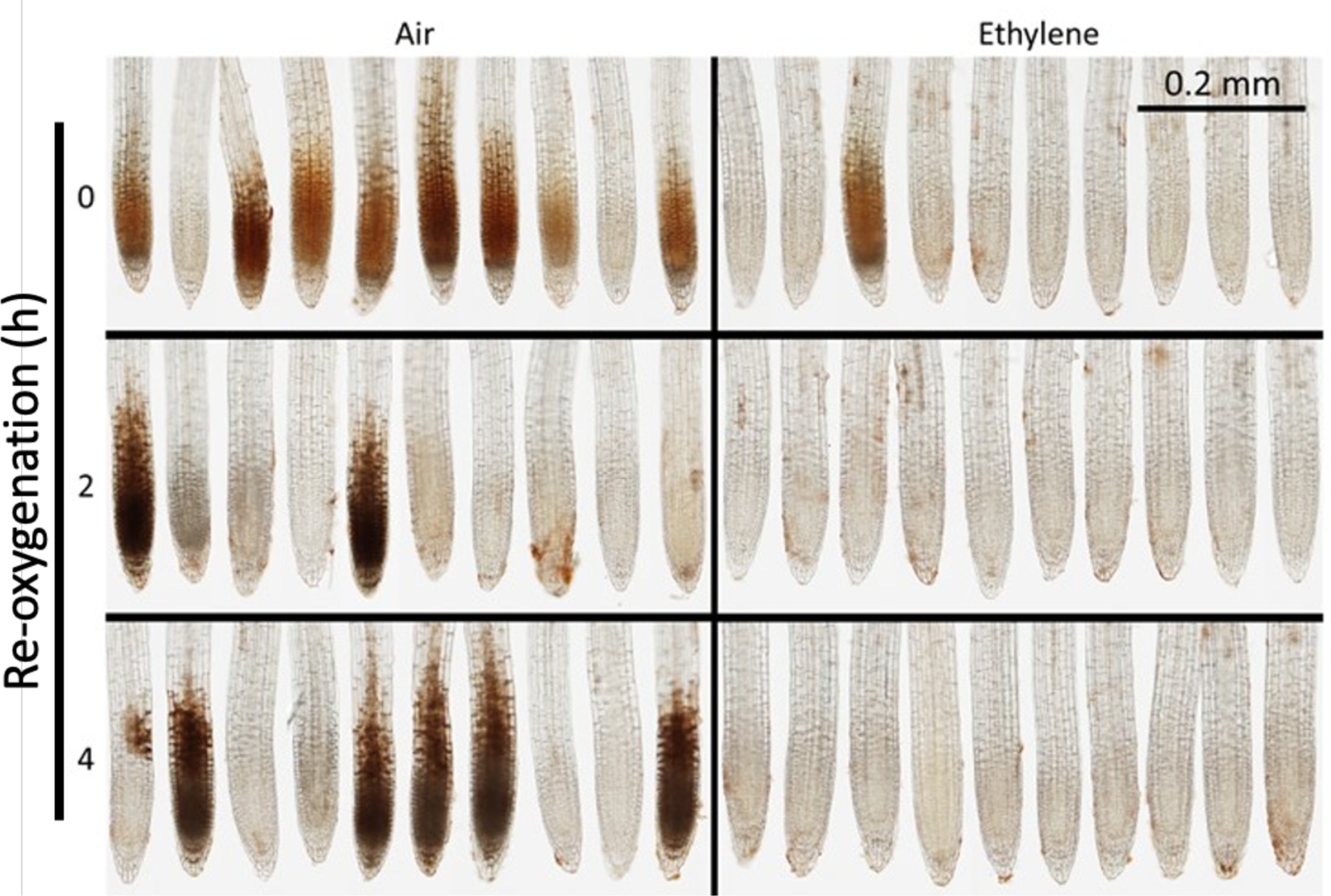
Ethylene limits H*_2_*O*_2_* accumulation during subsequent hypoxia and re-oxygenation. Representative DAB staining for H_2_O_2_ of 5-day old Arabidopsis seedling root tips after air or ethylene pre-treatment followed by 4 h of hypoxia and 2 and 4 h of re-oxygenation (n=10). Experiment was repeated twice with similar results.

**Supplemental Figure 9:**
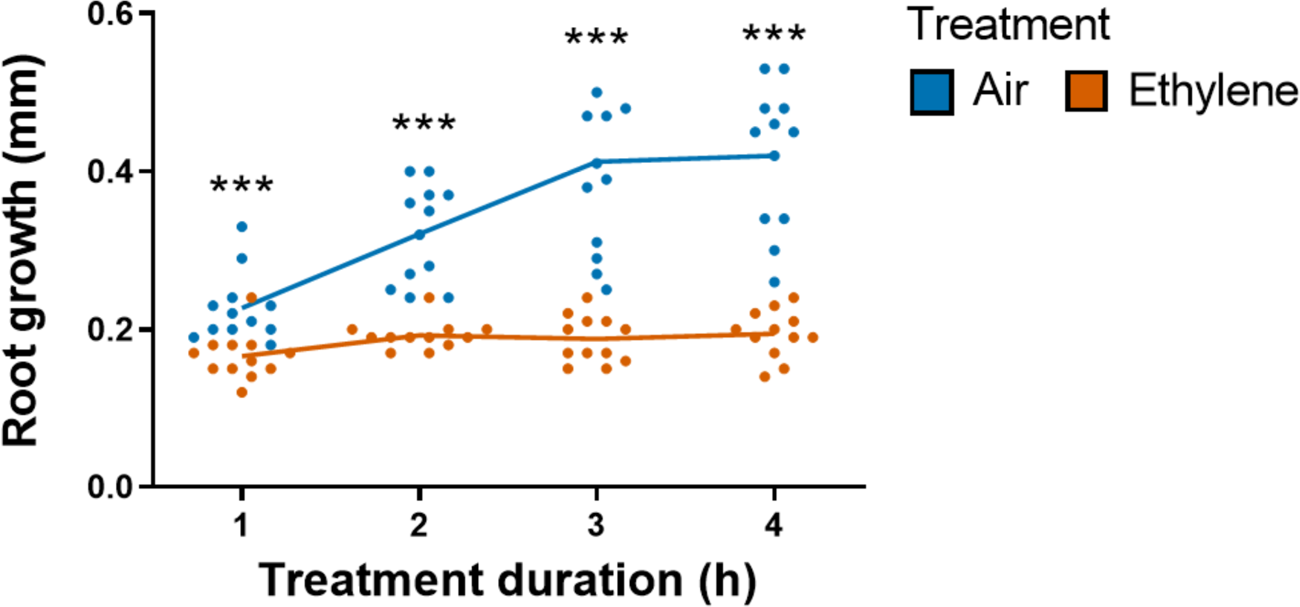
Ethylene rapidly slows down growth of primary roots. Root growth (increase in root length) of 4-day old Arabidopsis seedlings during 4h of air (blue) or ∼5 μL·L^-1^ ethylene (orange) treatment compared to root length at t=0. Asterisks indicate significant differences between air and ethylene per time point (***p<0.001, Student’s t-test, n= 12 rows containing 23 seedlings each).

## Notes

### Competing Interest Statement

The authors have declared no competing interest.

